# Human endogenous oxytocin and its neural correlates show adaptive responses to social touch based on recent social context

**DOI:** 10.1101/2021.04.08.438987

**Authors:** Linda Handlin, Giovanni Novembre, Heléne Lindholm, Robin Kämpe, Elisabeth Paul, India Morrison

## Abstract

Both oxytocin (OT) and touch are key mediators of social attachment. In rodents, tactile stimulation elicits endogenous release of OT, potentially facilitating attachment and other forms of prosocial behavior, yet the relationship between endogenous OT and neural modulation remains unexplored in humans. Using serial sampling of plasma hormone levels during functional neuroimaging across two successive social interactions, we show that contextual circumstances of social touch facilitate or inhibit not only current hormonal and brain responses, but also calibrate *later* responses. Namely, touch from a male to his female romantic partner enhanced subsequent OT release for touch from an unfamiliar stranger, yet OT responses to partner touch were dampened following stranger touch. Hypothalamus and dorsal raphe activation reflected plasma OT changes during the initial interaction. In the subsequent social interaction, time- and context-dependent OT changes modulated precuneus and parietal-temporal cortex pathways, including a region of medial prefrontal cortex that also covaried with plasma cortisol. These findings demonstrate that hormonal neuromodulation during successive human social interactions is adaptive to social context, and point to mechanisms that flexibly calibrate receptivity in social encounters.

## Introduction

A hug from a friend, a caress from a lover, the secure embrace of a parent: touch is a predominant sensory channel for bolstering human connection and emotional attachment. The neural mechanisms supporting this vital role of touch are not fully understood, but there is evidence from rodents that tactile stimulation in social interactions can act as a major trigger for endogenous release of the neuropeptide oxytocin (OT) (*1*,*2*), which has been implicated in social attachment (*3*,*4*). It has therefore been suggested that social tactile stimulation such as affectionate stroking may elicit endogenous OT release in adult humans (*5*–*8*).

In this perspective, afferent tactile stimulation initiates a cascade of modulatory responses in the brain, mediated by specific neural populations in the hypothalamus. It is well-established that magnocellular oxytocin neurons in the paraventricular nucleus (PVN) of the hypothalamus send projections to forebrain and cortical regions mediating olfactory and somatosensory signaling in the brain (*9*–*12*). For tactile stimulation, this signaling may rely on a recently-discovered population of parvocellular OT neurons in the rat PVN selective for particular forms of affiliative touch (*1*, *13*, *14*). OT released into the bloodstream via a PVN-pituitary pathway modulates the action of vasculature and smooth muscle (*15*), notably during processes surrounding reproductive behavior in both sexes, as well as parturition and lactation in females (*16*, *17*).

OT’s broad role in parental nurturance and affiliative behavior may reflect a functional extension of its influence on these core reproductive and maternal behaviors (*18*), many of which rely on sensory cues such as touch and olfaction, encompassing even cross-species interactions (*19*–*21*). Many OT-relevant sensory stimuli likely involve central pathways of OT in the brain, but peripheral mechanisms of release can also be triggered by stimulation of the genitals, the nipples, or the vagal nerve. However, despite evidence for afferent-driven central OT release, it is also acknowledged that OT-mediated social neuromodulation is highly dependent on factors such as social familiarity (*22*–*24*) and current physiological state (*25*,*26*),though this has been little explored for human endogenous OT. For example, endogenous OT can mediate prosocial allogrooming behavior in mice (*27*), but OT and OT receptor genotype have also been shown to play roles in antagonistic social behaviors such as defense of offspring (*28*) and aggression (*29*,*30*). Further, optogenetic manipulation of the same PVN OT neurons in freely-behaving mice can result in either prosocial or antagonistic behavior (*31*). These observations suggest that OT’s role in social behavior is multivalent and situational (*32*, *33*). Proposed functional roles for OT as selectively modulating affiliative social relationships (*34*) or maintaining allostatic stability (*25*) accommodate such differential, context-dependent effects. Nonetheless, a marked gap remains to be bridged between stimulus-driven and context-sensitive frameworks in charting the neural mechanisms of OT effects on social behavior.

In humans, it is not known whether endogenous OT changes and their neural correlates during social interactions may be selectively modulated by specific contextual conditions. However, the influence of exogenous, intranasal OT (IN-OT) administration on behavioral and neural outcomes has been widely studied with regard to social stimuli (*eg*, *35*,*36*). IN-OT has shown varying effects on social outcome measures such as face processing (*37*–*39*), empathy (*40*), romantic relationships and bonding (*41*–*43*), and romantic touch (*44*).

Neuroimaging studies of IN-OT manipulations indicate that many relevant changes occur at the cortical level (*45*,*46*), suggesting more complex modulatory pathways than is implied by a stimulus-driven model focused on afferent-subcortical signaling. However, as there is uncertainty surrounding the mechanisms of action of IN-OT and its degree of equivalence to endogenous release (*47*–*50*; but see *51*), it is crucial to investigate endogenous OT changes and their neural correlates in humans for a fuller understanding of the relevant mechanisms, as well as their potential limits and parameters.

In this study, we therefore combined serial sampling of plasma OT with functional magnetic resonance imaging (fMRI), to examine whether social interactions involving touch can evoke endogenous changes in plasma OT in human females. We predicted that touch from a socially familiar person (a romantic partner) would evoke greater endogenous OT changes than touch from an unfamiliar person (a nonthreatening stranger), thus allowing investigation of the neural responses associated with this modulation. We expected engagement of hypothalamus and other key regions associated with conserved circuitry of OT modulation and receptor expression in different species (such as amygdala, medial prefrontal cortex, and cingulate). The paradigm also allowed for investigation of the question of whether brain regions with differential responses to touch on arm as compared to palm skin would show OT-dependent modulation. This is because OT has been proposed to play a role in signaling of a specific subtype of C afferent nerve (C-tactile or CT), found in hair-follicle-containing skin and implicated in affective touch (*5*, *52*–*54*).

Crucially, we also explored whether any such changes would be modulated by recent social interaction history with familiar or unfamiliar others. To investigate this key temporal aspect of social context, we modeled changes in plasma OT alongside neural responses to partner and stranger touch across two successive interactions: one in which the stranger preceded the partner, and one in which the partner preceded the stranger. This allowed us to identify brain regions in which hemodynamic responses changed as a function of endogenous OT levels over the experimental session, revealing context-sensitive, OT-mediated engagement of both subcortical and cortical systems.

## Methods

### Participants

42 females in committed heterosexual romantic relationships of at least one year (mean age 24.6 years, SD=4.6, Table S1), participated in the study with their male partners (mean age 26.8 years, SD=6.0). Female participants were included if they were between 19-40 years, were not pregnant or breastfeeding, were free from estrogen-based contraceptive use, and were not undergoing current or recent hormone therapy. The female in the couple participated in fMRI scanning and provided plasma samples, while the male partner administered touch during the experiment.

This study took a female-first strategy (*55*), in order to limit any confounding sex specific effects and between-sex variability in OT and cortisol responses, and also in light of the predominance of males in human OT studies. We collected cycle phase self-report (with the aid of apps for most cycling participants) to rule out any variability which might be associated with relative estrogen increases during the late follicular phase, in light of evidence for OT-estrogen interactions in the context of sexual responses and parturition (eg *56*–*58*; but see *59*,*60* on the lack of evidence for effects of cycle-related variability on neural and physiological measures and gene expression in mice and humans).

Ethical approval was obtained from the Regional Ethical Review Board in Linköping, Sweden. All participants gave informed consent in accordance with the Declaration of Helsinki and were compensated at 400 SEK (~45 USD)/h.

### Procedure

An indwelling magnet-safe catheter was inserted into the cubital vein of the female participants’ left arm 40-60 min before the scanning session, to reduce the possibility of short-term effects of the challenge of needle insertion on plasma hormone levels during the main experiment. This took place in the same building as the MR suite.

In the ensuing 45-60 min the participant and her partner filled out two questionnaires (separately); the Couples Satisfaction Index (CSI; *62*) and the State-Trait Anxiety Inventory (STAI; *63*) and received instructions about the experiment (together). The participants and their partners were requested not to touch each other during this period. They were also informed of the presentation order (partner or stranger first), and were briefly introduced to the stranger, before going into the scanner. Before the partner/stranger entered the MR room for a functional run, the participant was informed about who entered the room over the audio system. This was to reduce any uncertainty as to the identity of the person delivering the touch, to avoid the risk of increasing psychological distress for the participant, as well as any potential increase in variance in the measures collected. Partners that were to deliver touch in the second run waited in a furnished staff break room until they were accompanied to the MR room.

### Experimental design and session structure

The experimental paradigm implemented a 2×2×2 factorial design: “familiarity” (partner/stranger, blocked by run), “order” (partner or stranger in the first encounter, counterbalanced), and “site” of the touch (arm or palm). Each run consisted of twelve 12-s touch stimulation trials, with arm and palm stimulation pseudorandomized within each of the two 7-min functional runs, and a jittered 21-30s intertrial interval. The first functional run was preceded by a T1-weighted anatomical scan. In the ~27m interval between the first and second functional runs, T2-weighted anatomical scans, resting state, and diffusion data (not analyzed here), were collected (Fig 1).

**Figure 1.**
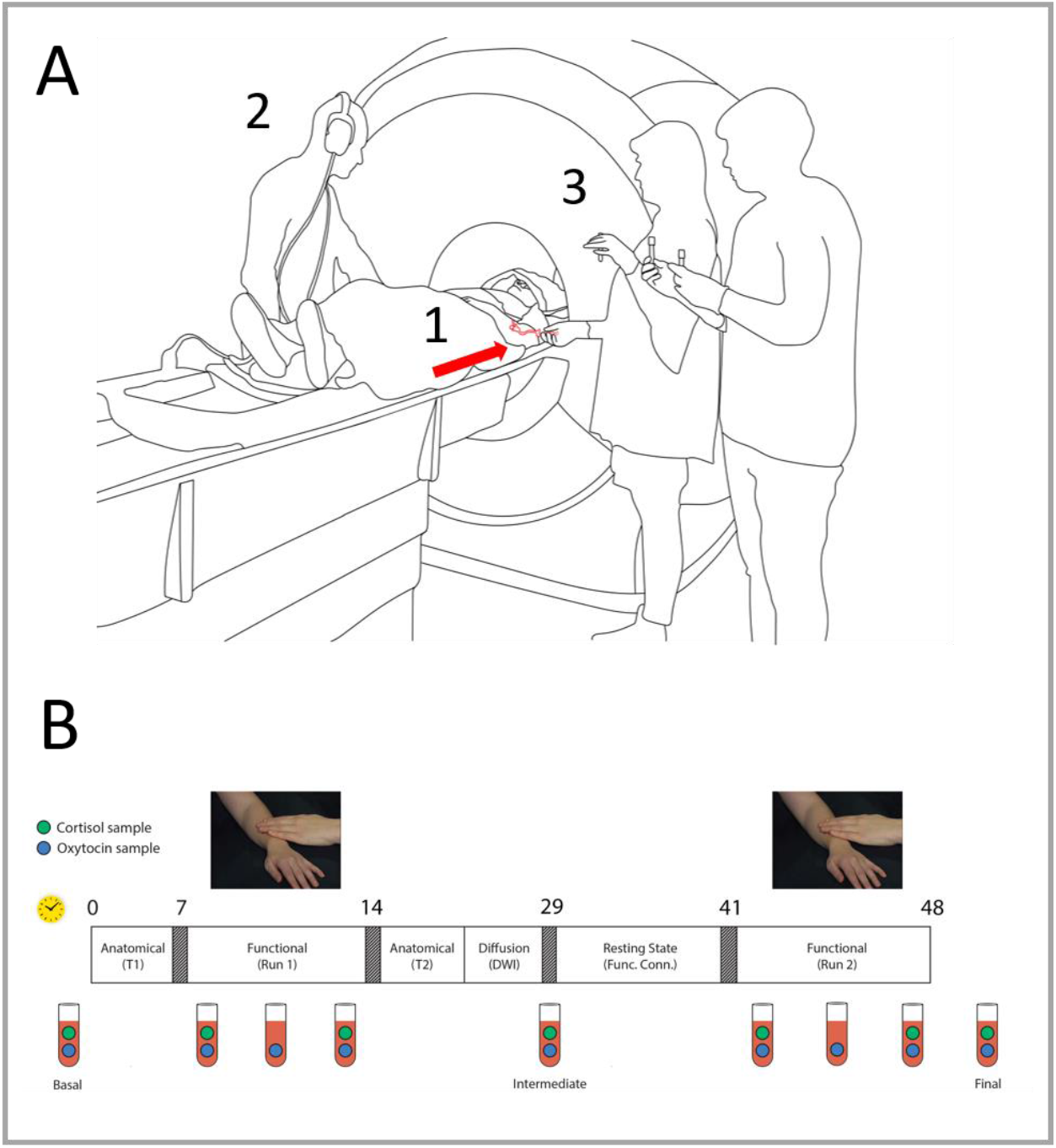
**A.** fMRI experiment setup. 1) Indwelling catheter in female participant’s left arm (arrow); 2) participant’s male partner or unfamiliar stranger caressed participant, following audio prompts; 3) serial blood samples were collected from the catheter. **B.** Structure of fMRI experimental session with serial sampling of plasma oxytocin and cortisol. Rectangle depicts time course of experiment, with approximate elapsed minutes shown above (yellow clock symbol). Two functional runs with partner and stranger touch, in counterbalanced order, were separated by ~27 min. 3 baseline OT samples (1,5,9) and 3 serial samples were collected for each run (2-4, 6-8). Clock symbol indicates time in minutes. Blue dots in the vial symbols indicate oxytocin samples, green dots indicate cortisol samples. The first functional run was preceded by acquisition of an anatomical (T1-weighted) image, while between functional runs additional anatomical and functional scans were acquired: T2-weighted anatomical image, diffusion-weighted imaged, and resting-state.

Touch stimulation was delivered manually by the male interactant (partner and stranger). Caressing strokes were delivered to the right dorsal arm and palm, with timing and touch site guided by auditory cues via headphones (Fig1A). The interactant was positioned beside the scanner bore on the right side of the participant. In the initial encounter (the first of the two functional runs), participants either received touch from their partner or the stranger, and vice-versa in the second encounter (N=24 and N=18, respectively). In the last 7s of each trial, the participant rated the pleasantness of the touch on a visual analogue scale (VAS), anchored with “most unpleasant imaginable touch” on one extreme and “most pleasant imaginable touch” on the other. Responses were made with the right hand using a response pad system (4-Button Diamond Fiber Optic Response Pad, Current Designs).

16 serial blood samples (of which 9 OT and 7 cortisol; < 70 ml total) were collected from each participant during the session: a pre-run OT and cortisol baseline for each of the two runs (preceding the T1 anatomical and preceding the resting state scans, respectively); three samples during each run at 1m, 3:30 min (OT only), and 6:30m; and a final post-session sample outside the scanner (Fig 1B). All blood samples were collected by a nurse and an assistant, who were positioned beside the scanner bore on the left side of the participant. Samples were collected through the indwelling catheter using vacutainer tubes (which rely on the vacuum action of puncturing of a vial’s rubber seal). The two functional runs were separated by ~27 min, allowing OT levels to return to baseline between runs. Samples collected in EDTA-tubes were used for OT analysis and samples collected in serum-gel tubes were used for cortisol analysis. All samples were centrifuged at 4°C at 10,000g for 10 min and plasma and serum were then aliquoted and stored at −20° C until analysis.

After the MR session, female participants evaluated how relaxing they had found partner and stranger touch, how attractive and trustworthy they had found the stranger, and how relaxing the interaction with the nurse had been. All ratings were performed using a visual analogue scale (VAS), with −10 as the most negative rating and +10 as the most positive rating.

### Hormone analysis

Plasma samples for OT analysis were extracted using acetonitrile precipitation (Merck Millipore: Human Neuropeptide Magnetic Bead Panel 96-Well Plate Assay Cat. # HNPMAG-35K) and OT concentrations were then determined using the Oxytocin ELISA kit (Enzo Life Sciences; sensitivity > 15.0 pg/ml, intra-assay precision 10.2-13.3 % CV, inter-assay precision 11.8-20.9 % CV).

Plasma cortisol levels were determined using the Cortisol Parameter Assay Kit according to the manufacturer’s recommendations (R&D Systems, Minneapolis, Minnesota, USA) (sensitivity 0.071 ng/mL, precision 10.4%). Cortisol analysis serum samples were diluted 60 times preceding analysis. Pretreatment steps of the serum samples resulted in a dilution factor of three and the pretreated serum samples required an additional 20-fold dilution in Calibrator Diluent RD5-43.

For both the OT and cortisol analyses, standards and controls were implemented according to manufacturer recommendations. Washing procedures were performed using a Wellwash™ Microplate Washer (ThermoFisher Scientific, Waltham, Massachusetts, USA) and the absorbance was read using a Multiskan™ FC Microplate Photometer (ThermoFisher Scientific, Waltham, Massachusetts, USA). The color development of the samples was read for OT at 405 nm (background correction at 571 nm) and for cortisol at 450 nm (background correction at 571 nm). SkanIt™ Software was used for creation of standard curves, curve fitting and calculation of concentrations (ThermoFisher Scientific, Waltham, Massachusetts, USA).

### fMRI data acquisition

fMRI data were acquired using a 3.0 Tesla Siemens scanner (Prisma, Siemens) with a 64-channel head coil. For each functional run, 456 2D T2*-weighted echo-planar images (EPIs) were acquired (repetition time: 901 ms; echo time: 30 ms; slice thickness: 3 mm; no slice gap; matrix size: 64 × 64; field of view: 488 × 488 mm^2^; in-plane voxel resolution: 3 mm^2^; flip angle: Ernst angle (59°). Three dummy volumes were acquired before each scan to ensure that data collection started after magnetizations reached a steady state.

### fMRI preprocessing and analysis

Preprocessing and statistical analysis of MRI data were performed using Analysis of Functional Neuroimages (AFNI) statistical software (version 19.1.12). Functional data were first de-spiked. Each EPI volume and the T1 were then aligned to the EPI volume with the minimum outlier fraction (using the AFNI outlier definition) to correct for motion. Functional images were warped to the MNI 152 template using a combination of affine and non-linear transformations (*63*). Finally, spatial smoothing was applied with a 10 mm full-width at half-maximum filter. Residual effects of head motion were corrected by including the estimated motion parameters (and their derivatives) as regressors of no interest. A motion censoring threshold of 0.2 mm per TR was implemented in combination with an outlier fraction threshold of 0.1. Volumes violating either of these thresholds were subsequently ignored in the time-series regression.

For each participant, whole-brain voxel-wise general linear models (GLM) were created for each of the two runs using 3dDeconvolve. One regressor (convolved with a standard model of the hemodynamic response function, HRF) modeled each of the conditions: partner arm, partner palm, stranger arm, stranger palm. To determine specific effects of partner and stranger touch regardless of stimulation site, GLMs were created with partner touch and stranger touch as regressors. Each regressor modeled 10 seconds within the 12s touch interval, beginning two seconds after the onset of touch stimulation and ending with stimulation offset.

We also included additional regressors of no interest to model the effects of motor responses during the rating of the touch stimuli.

At the group level, the AFNI program 3dClustSim was used to determine cluster-size thresholds for identifying effects significant at *α*= 0.05 family-wise-error (FWE) corrected (within-cluster). Average spatial smoothness estimates, across all participants, used by 3dClustSim were obtained using the 3dFWHMx function with the ACF flag, as per current AFNI recommendations (*63*). For each analysis, we report brain areas which were activated at a voxel-wise p-value thresholds of *p* = 0.002 (activations present at p = 0.005 are included in Supplementary Tables. Since AFNI outputs a single peak coordinate for each surviving cluster, a custom script was used to extract the coordinates for the first 10 peaks with the highest T scores for each cluster.

## Results

### Hormone analyses

#### OT levels

Oxytocin levels measured before and during each touch session were entered into a mixed linear model using SPSS version 27, n = 27. Familiarity (partner or stranger) and sample timepoint (pre-run baseline and three samples for each functional run) were included as within-subject factors, whereas order (partner or stranger first) was included as a between-subject factor. Since the Intraclass Correlation Coefficient (ICC) was high (0.73), both fixed and random intercepts were included in the model, and marginal means were estimated through maximum likelihood method. Post-hoc pairwise comparisons were performed to investigate significant interactions, applying Bonferroni correction to adjust for multiple comparisons.

The linear mixed model revealed a three-way interaction between the factors familiarity, order and sample timepoint (F_(3, 183.180)_ =3.034, p=0.031) as well as a two-way interaction between the factors familiarity and order (F_(1, 183.169)_ =11.216, p=0.001). Post-hoc t-tests revealed that the contribution of timepoint to the three-way interaction was driven by a difference between the first and middle samples in the functional run during stranger touch in the partner first group only (p=0.027). Marginally below-alpha differences were seen between middle and final samples during the run, pre-run baseline and the first sample during the first run in the partner first group (p=0.054 and p=0.070, respectively). There were no significant main effects, despite a trend for the main effect of order (F_(1, 27.087)_ =3.855, p=0.060).

### Cortisol levels

Student’s T-tests were first performed to test for differences in basal cortisol levels depending on time of day (i.e. sessions beginning at 9:00, 12:00 or 15:00). Time of day did not have a significant effect on the participant’s basal cortisol levels.

Cortisol levels measured before and during each touch session were entered into a mixed linear model using SPSS version 27 (n = 26 with complete sample series). As with the oxytocin analysis, familiarity (partner or stranger) and sample timepoint (pre-run baseline and two samples for each functional run) were included as within-subject factors, whereas order (partner or stranger first) was included as a between-subject factor. Since the ICC was high (0.81) both fixed and random intercepts were included in the model and marginal means were estimated through maximum likelihood method. Post-hoc pairwise comparisons were performed to investigate significant interactions, applying Bonferroni correction to adjust for multiple comparisons.

The linear mixed model revealed an interaction between the factors familiarity and order (F=54.89_1,25_, *p* <0.001). This interaction was driven by higher cortisol levels for participants for whom the stranger was presented in the first run, compared to those for whom he was presented second (*p* = 0.008). The model also revealed significant main effects of familiarity (F=15.67_1,25_, *p* <0.001) and sample timepoint (F=3.16_1,25_, p=0.045). Post-hoc t-tests revealed that cortisol levels were higher in the beginning of the touch session compared to the end (*p* = 0.039) and that stranger touch elicited a greater cortisol increase compared to partner touch (p<0.001; Fig 3A).

#### Oxytocin x cortisol interaction

To investigate whether and how OT and cortisol levels interacted, we defined an additional mixed linear model in SPSS version 27 (n = 26). In addition to familiarity (partner or stranger) and sample timepoint (pre-run baseline and three samples for each functional run), a within-subject factor for hormone (OT or cortisol) was also included, and order (partner or stranger first) was included as a between-subject factor. Considered that ICC values were high for both OT and cortisol individual models, and we did not assume independence between the two hormone values for each participant, both fixed and random intercepts were included, and marginal means were estimated through maximum likelihood method. Post-hoc pairwise comparisons were performed to investigate significant interactions, applying Bonferroni correction to adjust for multiple comparisons.

An interaction between the factors order and hormone (F_1,180.593_ =28,751, p<0.001) was driven by higher OT but lower cortisol levels in partner first compared to stranger first (OT Mean±SEM: partner first: 67.781±6.019, stranger first: 42.959±1.816, p=0.017; Cortisol Mean±SEM: partner first: 80.346±5.486, stranger first: 101.633±6.306, p=0.040). There was also a main effect of hormone (F_1, 180.593_=68.574, p<0.001), with higher overall cortisol levels than OT levels, as well as an interaction between familiarity and order (F_1, 178.355_=10.565, p=0.001; also found in the two previous individual models).

### Behavioral measures

#### Ratings

Partner touch was rated as more pleasant than stranger touch (F_1, 33_=30.032, p < 0.001) with a main effect of higher ratings for arm (F_1, 33_=11.070, p = 0.002, effect size f = 0.7, partial η2 = 0.33 at power (1-β error probability) = 0.8, α = 0.05). A significant three-way interaction (F_1, 33_=4.730, p = 0.037) was driven by lower ratings for stranger touch on the palm compared to stranger touch on the arm, for participants who received stranger touch first, (t=16, p = 0.007, d = 1.99).

The stranger was rated as positively trustworthy, with a mean rating of 3.96 on a visual analogue scale from −10 (untrustworthy) to 10 (trustworthy). Stranger attractiveness was near a neutral midpoint, with a mean rating of 0.95 on a visual analogue scale from −10 (unattractive) to 10 (attractive). For participants starting with partner touch, evaluation of relaxation for partner touch correlated with how trustworthy participants rated the stranger (r = .77, *p* = 0.002). The interaction with the nurse was rated as non-stressful, with a mean rating of 5.28 on a scale of −10 (stressful) to 10 (calming). For participants in the stranger first group, the smaller the difference between pleasantness ratings for stranger and partner touch during the imaging session, the more relaxing participants rated stranger touch afterwards (r = −.70, P = 0.004). Within-session pleasantness ratings also predicted post-session evaluations of relaxation for partner and stranger touch independently (partner r = 0.83, *p* = 0.0002; stranger r = 0.80, *p* = 0.001).

#### Self-report questionnaires

CSI scores indicated that both female and male participants were satisfied with their relationships (females: mean=139.8, sd=17.2, males: mean=139.6, sd=15.2). The total score is the sum of responses across all 32 items and can range from 0 −161. Scores below 104.5 suggest relationship dissatisfaction, while higher scores suggest greater levels of relationship satisfaction. Participants’ assessments of relationship quality correlated with their partners’ (r=0.57, p=0.0003). State-Trait Anxiety Inventory (STAI; *68*) scores indicated that no participants demonstrated clinically significant symptoms of anxiety (females STAI-S mean=35.6, sd=9.7; female STAI-T mean=40.3, sd=9.8; males STAI-S mean=31.6, sd=5.7; males STAI-T: mean=36.2, sd=6.7). Participants’ trait anxiety correlated inversely with the both the participant’s and her partner’s assessment of relationship quality (r=-0.40, p=0.01 for participant, r=-0.39, p=0.01 for partner). There were no other significant correlations among CSI, STAI-T, STAI-S, and VAS pleasantness scores (all ps< 0.01). Missing data in both the CSI and STAI questionnaires (maximum two missing values) was handled with hot deck imputation (*64*).

#### Hormonal cycles and OT levels

Of the 27 participants with full sample series included in the OT hormone analysis, 10 were not cycling (ovulation suppression by non-estrogen-based contraceptives, for example copper intrauterine devices), 13 were cycling naturally (using condoms as contraception, for example), and 4 did not report their cycle details. The mean OT of cycling participants fell within 1 standard deviation of noncycling participants for both partner and stranger (noncycling: 68.8 pg/ml ± 43.4 for partner, 64.1 ± 40.3 for stranger; cycling: 62.9 pg/ml ± 31.1 for partner, 48.6 ± 21.3 for stranger), indicating that cycle did not affect the variability of mean plasma OT. This was also the case for cycle phase among the cycling participants, of whom 4 were in the luteal phase (relatively stable but lower estrogen), 2 in the follicular phase (2 early, 2 in the later days associated with sharply increased estrogen preceding ovulation), and 7 were menstruating (decreasing to stable estrogen).

### Functional neuroimaging

#### Linear mixed-effects modeling with OT covariate

In order to reveal BOLD activations related to the order of the touch interactions (partner first, stranger first), the familiarity of the interactant (partner, stranger) and peak OT plasma levels, we performed a linear mixed-effects modeling analysis (3dLME in AFNI; Table s2). 3dLME was implemented because the analysis involved a between-subject factor (order), a within-subject factor (familiarity), and a quantitative variable or covariate (OT), modeled with random intercept and random slope (*70*). All analyses were thresholded at p = 0.002 as per current AFNI recommendations (*73*).

Since OT response was influenced by the order factor, the peak OT values (maximum percent change from baseline) for each participant (n=26 complete datasets) were centered around the mean of each of the 4 conditions (partner first, stranger first, partner second, stranger second) prior to the analysis. The 3dLME analysis revealed a 3-way interaction between familiarity, order, and OT in the right superior occipital gyrus, right cuneus, and right TP. Familiarity and OT interacted in right supramarginal/angular gyri (SMG/AG), right superior temporal gyrus (STG), right middle temporal gyrus (MTG), right temporal pole (TP), bilateral anterior cingulate cortex (ACC), and right medial prefrontal cortex (mPFC, on mid-orbital gyrus).

To further compare BOLD activity across familiarity and order factors with OT as a covariate of interest, the following t-tests were performed: independent t-tests between partner first and stranger first and partner second and stranger second; and paired t-tests between partner first and stranger second and between stranger first and partner second (Table S3).

##### Partner First vs. Stranger First

Hypothalamus and Raphe nuclei showed BOLD increases during partner touch in the first run. To explore whether these BOLD signal changes in hypothalamus covaried with OT levels, we confirmed whether all subjects included in the whole-brain analysis had representative data in those voxels to be included in a region of interest (ROI) approach. We used the Neurosynth database’s ‘association test map’ for the search term “hypothalamus” (https://neurosynth.org/analyses/terms/hypothalamus). This 1565-voxel map also encompassed regions outside the hypothalamus (e.g., thalamus and brainstem), so we used a restricted threshold to include only voxels located in the hypothalamus. The size of the final ROI was 111 voxels, and this was used in a subsequent SVC analysis including 25 of the initial 27 participants. Two participants (one per group) were excluded as they had data for less than 50% of the ROI voxels in at least one of the two functional runs (partner touch, stranger touch). Within this cluster, a subset of 15 voxels showing greater increases for partner touch than stranger touch survived stricter cluster correction at p = 0.002.

##### Partner Second vs. Stranger Second

BOLD increased for partner touch compared to stranger touch in parietotemporal clusters also seen in the interaction maps for familiarity x order x OT (right TP) and order x OT (right AG/SMG, right MTG, right TP, right STG), as well as additional clusters in right superior medial gyrus, bilateral midorbital gyrus and right superior frontal gyrus. These activations reflected a positive relationship between BOLD and the OT covariate, whereas this relationship was negative during subsequent stranger touch in the second run. The opposite was observed for participants for whom the stranger was presented in the first run.

##### Partner First vs. Stranger Second

There was no activation difference between partner and stranger touch in the partner first group. *Stranger First vs. Partner Second:* There was no activation difference between partner and stranger touch in the stranger first group.

#### OT regressor

To discover activation corresponding with the overall temporal pattern of the endogenous OT response over the sample series in each functional run, we used each participant’s serial plasma OT levels to create a custom regressor for each individual. Points between the multiple samples were linearly interpolated and the resulting function was convolved with the canonical hemodynamic response function (HRF). N=23 participants had complete data series for both functional runs (Fig S1). Adding exploratory time lags of 1, 1.5, 2, 2.5 and 3 min with this regressor allowed identification of BOLD activation which *preceded* the pattern of serial plasma OT level changes in these five time windows. Thus, it captured potential central modulation corresponding to the peripheral changes in plasma OT observed after various delays.

This analysis revealed brain areas showing significant interactions between familiarity and order at both 2 and 2.5 minutes, with higher BOLD during stranger first than partner first, but no difference between stranger second and partner second. No activations were observed for the remaining time lags. *2 min*. Bilateral precuneus, right SMG, left inferior parietal lobule, right posterior cingulate cortex, right postcentral gyrus, and right parietal operculum showed increased BOLD corresponding to the pattern of changes in OT levels observed 2 minutes after the activation (p = 0.002; Table S4). *2.5 min*. Two and half minutes before observed changes in OT levels, increased BOLD in bilateral precuneus and right paracentral lobule corresponded with the temporal pattern of OT changes (all at p = 0.002, Table S4).

### Cortisol analyses

#### Linear mixed model with cortisol covariate

As for OT, we searched for brain correlates of touch interactions as influenced by familiarity (partner, stranger), order (partner first, stranger first), and cortisol plasma level, performing a linear mixed-effects modeling analysis (3dLME in AFNI; n=26). Again, we adopted the model with random intercept and random slope and cortisol mean values (percent change from baseline) for each participant were centered around the mean of each of the 4 conditions (partner first, stranger first, partner second, stranger second) prior to the analysis. Results did not show any significant main effect or interaction.

#### T-tests with Cortisol covariate

As for oxytocin, in order to compare neural activity across the toucher and order factors with cortisol as covariate of interest, we ran the following t-tests: paired t-tests between “partner first” and “stranger second” and between “stranger first” and “partner second”; independent t-tests between “partner first” and “stranger first” and “partner second” and “stranger second” (Table s7).

##### Partner first vs. stranger first

In the first run, the two groups did not differ in terms of activation according to the identity of the stroker.

##### Partner second vs. stranger second

In the second run, there was no differential activation between partner first and stranger first groups related to the identity of the stroker.

##### Partner first vs. stranger second

For participants who had partner touch first, partner compared to stranger touch covaried with cortisol in the left anterior cingulate cortex, right superior medial gyrus, bilateral ventromedial prefrontal cortex, bilateral calcarine gyrus, right lingual gyrus, left temporal pole and left parietal operculum (all at p = 0.002), and in the bilateral posterior cingulate cortex at p = 0.005.

##### Stranger first vs. partner second

When comparing brain activity in the two runs in the group of participants who had stranger touch first, we found no cortisol-related differences in brain activity between partner and stranger touch.

### Oxytocin-independent analyses

#### General linear model

To examine BOLD responses to the touch conditions independently of OT or cortisol levels, an ANOVA was performed (using the 3dANOVA program in AFNI) with the following two factors: familiarity (partner, stranger) and site (arm, palm). The following analyses include participants for whom complete fMRI data was collected, even those without a complete series of plasma hormone samples (n = 35). The remaining 7 participants had partial data sets due to technical issues during data collection or elective withdrawal from the experiment (see s1).

##### Familiarity x site interaction

The two factors interacted in the left postcentral and precentral gyri (p = 0.002), ipsilateral to stimulation. We conducted a repeated-measure ANOVA from betas extracted from these two clusters. Results showed that the interaction can be explained in terms of higher activation for stranger than partner for palm but not for arm for both the left (ipsilateral) postcentral gyrus (palm: t(34) = 3.188, p = 0.003; arm t(34) = - 1.863, p = 0.071) and right (ipsilateral) cerebellum (palm:: t(34) = 2.364, p = 0.024; arm t(34) = −2.097, p = 0.044). *Main effect of site*. Palm touch showed higher activation than arm touch in bilateral postcentral and precentral gyri, bilateral supplementary motor area (SMA), left superior parietal lobule, bilateral superior frontal gyrus, right parietal operculum, right SMG, bilateral cerebellum and right middle frontal gyrus. The reverse contrast (arm > palm) showed greater activation in the left superior temporal gyrus, left postcentral gyrus, left precuneus, and left posterior insula/parietal operculum (35 voxels, not surviving cluster-threshold correction; Table s6). *Main effect of familiarity*. Touch delivered by the stranger resulted in higher activation of the right middle frontal gyrus and the left inferior frontal gyrus compared to partner touch.

#### Conjunctions

To identify common activation across all 4 conditions, we created single condition maps by contrasting each condition against baseline activity at p < 0.002 (Table S5). We then calculated conjunction maps by considering all common suprathreshold voxels for the four conditions (partner first ⋂ stranger first ⋂ partner second ⋂ stranger second), as well as intersections between the initial encounter and second encounter only (partner ⋂ stranger, for each run). We also created within-subject conjunction maps between the two runs for participants receiving partner touch in the first encounter (partner first ⋂ stranger second,) and those receiving stranger touch in the first encounter (stranger first ⋂ partner second). Finally, we tested for conjunctions across both partner and stranger conditions.

##### Partner first ⋂ stranger first ⋂ partner second ⋂ stranger second

Brain areas showing increased activity irrespective of condition were: bilateral supramarginal gyrus (SMG), bilateral postcentral gyrus, left precentral gyrus, left superior parietal lobule (SPL), left parietal operculum (PO), bilateral supplementary motor area (SMA), right inferior frontal gyrus (IFG), bilateral anterior insula, right cerebellum, right supplementary motor area (SMA), and right amygdala. There was also a common deactivation in the right precentral gyrus. A complete list of conjunctions can be found in the Supplementary Materials.

## Discussion

A single social interaction with a familiar partner is one instance of many over the course of the relationship. On the other hand, today’s familiar friend was yesterday’s stranger: an interaction with a person one has never met can lay the groundwork for future interactions. Neural and hormonal mechanisms underlying such responses during successive social interactions must therefore not only be stable (*25*), but also flexible, depending on social context and history. The present findings shed light on the potential contribution of OT to adaptable neuromodulation during social encounters with both familiar and unfamiliar individuals. They demonstrate that touch-mediated social interactions in human females can influence endogenous OT and brain responses in a covariant manner. Moreover, these hormonal and neural changes are modulated by the familiarity of the person delivering touch and the recent history of social interaction.

The influence of these factors on within-subject endogenous OT changes was reflected in an interaction between the familiarity of the social interactant (partner or stranger) and the order of his presentation (partner first or stranger first). Here, OT modulation was driven by a greater increase in plasma OT responses for the stranger following partner touch, whereas there was no corresponding increase for partner touch following stranger touch (Fig 2A). This dependence of OT responses on both familiarity and presentation order is consistent with evidence for cumulative effects of central OT exposure over repeated interactions with specific individuals (*27*), as well as the established context-sensitivity of OT effects in nonhuman mammals (*26*). Yet beyond this, social familiarity and presentation order also interacted with a series timepoint among the eight plasma samples collected during the experimental session. This was driven by an initial dip and a subsequent recovery to an above-baseline level during touch from the unfamiliar stranger—but only when he was preceded by the familiar partner—suggestive of a context-dependent adjustment in the OT response.

**Fig 2.**
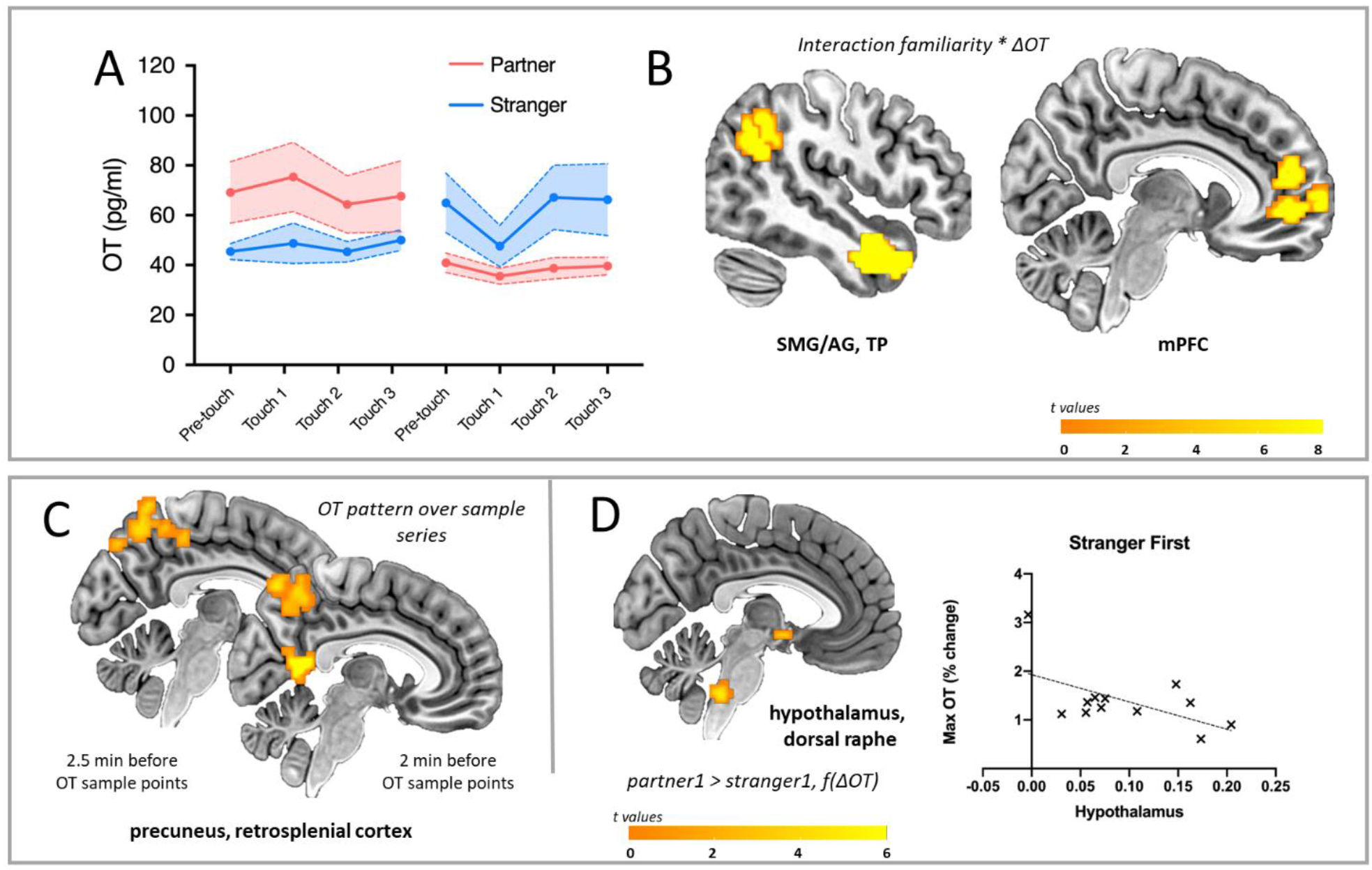
Endogenous oxytocin (OT) changes and covariant brain responses. **A.** Familiarity, order, and sample timepoint influenced plasma OT levels: familiarity, order and sample timepoint interacted (F_(3, 183.180)_ =3.034, p=0.031) as did familiarity and order (F_(1, 183.169)_ =11.2 1 6, p=0.001), with increased OT when the partner was the interactant in the first encounter. The contribution of sample timepoint lay in a dip and recovery during stranger touch only when preceded by partner touch (p=0.027). **B.** Familiarity and % OT change interacted in BOLD modulation in parietotemporal clusters along right SMG/AG, TP, mPFC (pictured), as well as right STG, MTG, and bilateral ACC (3dLME model, p < 0.002). **C.** The temporal pattern of OT-BOLD changes in bilateral precuneus preceded sampling by 2.5 min, with retrosplenial cortex also emerging at 2 min, with greater OT-BOLD covariance for stranger touch during the initial encounter. **D.** OT-BOLD covariance in hypothalamus and dorsal raphe (not shown) were driven by a greater decrease for stranger touch during the initial encounter (scatterplot shows beta values unweighted by OT covariate). *BOLD = blood-oxygen-level-dependent, 3dLME = 3-dimensional linear mixed effects, SMG/AG = supramarginal gyrus/angular gyrus, TP = temporal pole, mPFC = medial prefrontal cortex, STG = superior temporal gyrus, MTG = middle temporal gyrus, ACC = anterior cingulate cortex, f(ΔOT) = as a function of the change in OT. All maps thresholded at p < 0.002, corrected*.

### BOLD correlates of endogenous OT changes

The interactions observed in OT levels between social familiarity and partner/stranger presentation order were mirrored in OT-covariant BOLD responses in key partietotemporal and frontal regions, on a whole-brain level (Fig B,C,D). In particular, OT change interacted with both familiarity and presentation order in a temporal pole (TP) cluster; in SMG/AG, IFG, mPFC, and superior frontal gyrus, OT changes interacted with the familiarity of the individual delivering the touch. These partietotemporal and medial prefrontal regions have been implicated in individual intranasal-OT studies of partner-stranger interactions (*66*–*69*), animate visual social stimuli (*70*), as well as in meta-analyses of intranasal-OT fMRI activation (*71*,*72*) and even in tactile foot massage (*6*). They may thus reflect updating of contextual information, possibly enhancing the salience of incoming sensory signals (*73*–*75*) following the initial encounter. It is noteworthy that the relationship between BOLD and OT change in these regions was an inverse one, manifesting in greater BOLD increase alongside a relative decrease in OT change for partner vs stranger during the second encounter, further suggestive of a dynamic relationship between contextual factors and OT neuromodulation. Such a relationship may involve selective tuning and gain control of hormonal neuromodulation, more akin to a dimmer switch than an on-off button.

In the initial touch encounter, the preferential increase in plasma OT levels for the partner is consistent with familiarity-dependent effects of OT in rodents (*27*). This basic partner-stranger difference was reflected in hypothalamus OT-covariance, as a function of individual changes in plasma OT levels. This is in accord with the conserved mammalian neuroanatomy of central OT release, in which hypothalamic nuclei, particularly the PVN, synthesize OT and stimulate its release in the brain (*9*–*14*). In rats, stroking touch increases Fos protein expression in PVN (*52*), and recent evidence from freely-interacting female rats indicates that a population of parvocellular OT neurons in PVN is selectively tuned to social touch stimulation (*1*, *15*). In the present study, BOLD changes in dorsal raphe nuclei also covaried with plasma OT (Fig 2D), possibly reflecting descending influence in modulating touch signaling from the periphery. BOLD in both hypothalamus and dorsal raphe activation also showed a negative relationship with mean OT when the stranger touched first (Fig 3D), again suggestive of dynamic co-modulation.

**Fig 3.**
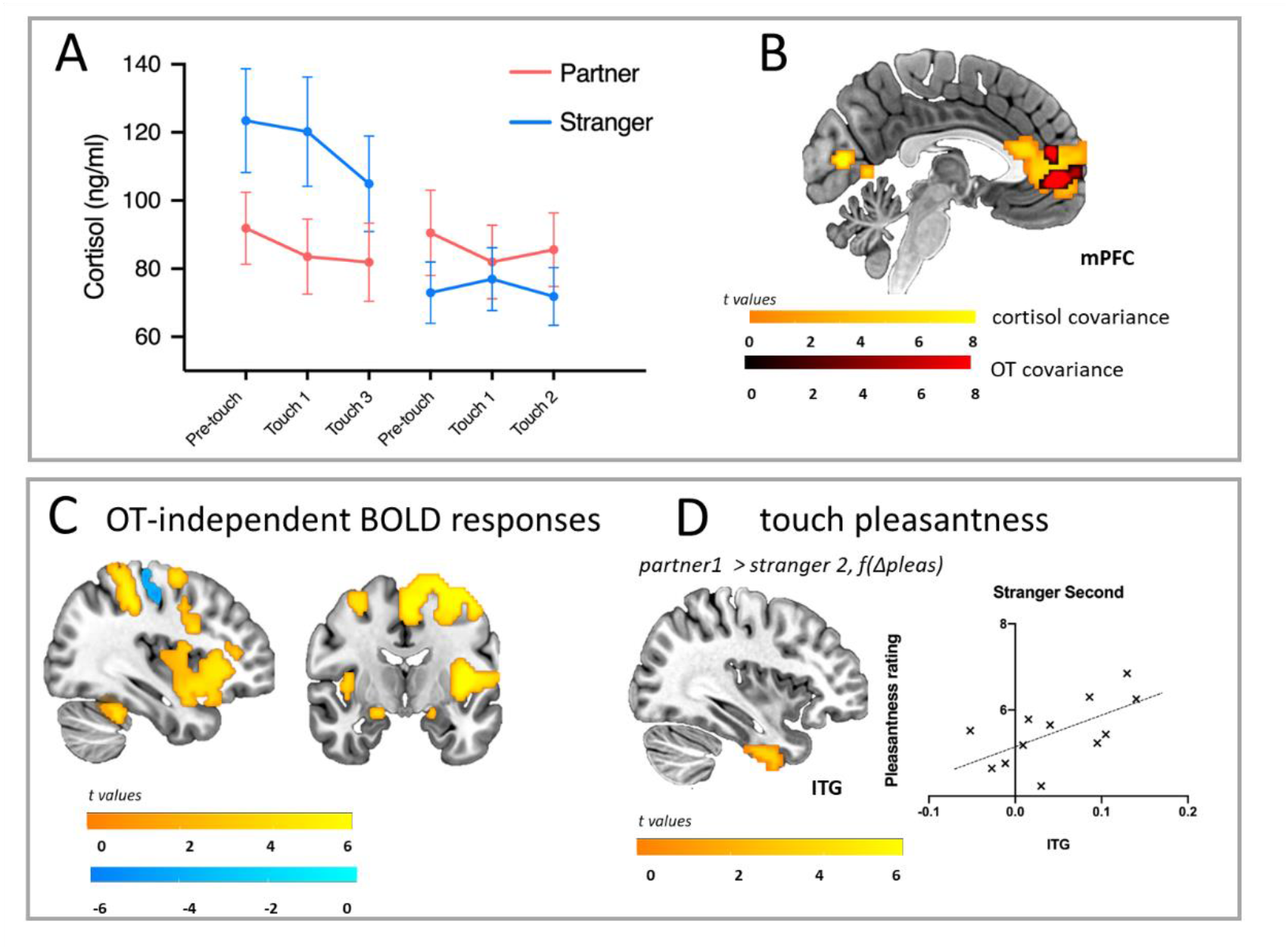
Plasma cortisol and covariant brain responses, and OT-independent brain activation. **A**. Familiarity and order interacted in plasma cortisol levels (F=54.89, *p* <0.001), with stranger touch eliciting a greater cortisol increase compared to partner touch, reflected in main effect of familiarity (F=15.67, *p* <0.001). There was also a main effect of sample timepoint (F=3.13, p=0.045), with levels generally declining over the session. **B.** BOLD signal change in regions including mPFC covaried as a function of cortisol levels, with partner > stranger (P <0.002), and a subset of mPFC voxels covarying with both OT (partner > stranger, second encounter) and cortisol (partner > stranger, initial encounter). **C.** BOLD changes in somatosensory and insular cortices, as well as bilateral amygdalae, across all touch conditions, independently of familiarity of person delivering touch, order, and OT levels (all Ps <0.002). **D.** ITG was sensitive to differences in partner and stranger touch pleasantness ratings (Δpleas), reflected in decreased BOLD covariance in ITG (*p* < 0.005) during stranger touch in the second encounter (mean pleasantness partner > stranger, *p* < 0.001). *BOLD = blood-oxygen-level-dependent, ITG = inferior temporal gyrus, mPFC = medial prefrontal cortex. All maps thresholded at p < 0.002, corrected*.

Central effects of IN-OT have consistently been found ~45 min post-administration (*75*–*77*), but there is less direct evidence about the timecourse of central endogenous OT release into the periphery in humans. We therefore developed an exploratory regressor based on the serial pattern of individuals’ OT levels. This allowed us to look “backwards” from the temporal pattern of the plasma OT sample series to any preceding hemodynamic activation that covaried with it. The pattern-covariant engagement of the precuneus at 2.5 minutes preceding sampling, and of precuneus, retrosplenial cortex, and mPFC at 2 minutes, is within the frame of the half-life of OT in blood (*78*) and may reflect events surrounding central OT release.

Retrosplenial cortex projects to mPFC (*79*), while precuneus is functionally connected to the SMG/AG region activated for partner during the second encounter. IN-OT administration can also induce changes in functional connectivity between precuneus and AG (*80*). These findings imply that precuneus, retrosplenial cortex, and mPFC may act as arbiters of activation in parietotemporal and limbic networks, potentially influencing responses to social touch via contextual integration and affect regulation processes. These OT-dependent neural dynamics may play a critical role in calibrating social receptivity, especially over multiple social encounters. It is not possible to determine the directness or direction of influence from these exploratory findings, but the temporal directionality suggests that cortical processing precedes plasma changes.

OT has been also implicated in stress regulation via corticotropin-releasing-hormone (CRH)-cortisol pathways and may act as a physiological regulator of acute stress-related responses (*81*–*85*). Here, mean plasma cortisol levels were higher for stranger than for partner during the first encounter (Fig 3A) and interacted with mean OT levels, indicating an inverse relationship between OT and (CRH)-cortisol in the circulation. Plasma cortisol levels covaried with BOLD for partner in the initial encounter in several regions, including a mPFC cluster which contained a subset of OT-sensitive voxels (Fig 3B). Further, both mPFC, implicated in cortical-amygdala signaling following IN-OT, and superior temporal gyrus (STG), implicated across a range of multisensory integration, including touch (*86*–*88*) decreased for initial stranger touch as a function of cortisol, compared to partner.

### Hormone-independent BOLD changes

BOLD responses in several key regions were independent of plasma OT or cortisol changes (Fig 3C), suggesting an absence of direct, covariant modulation by endogenous OT. The bilateral amygdalae were activated in a general fashion across all encounters (Fig 3C; Table S5), with left amygdala selective for stranger touch, particularly in the first encounter. Amygdala activation has been widely implicated in human IN-OT studies (*89*–*92*), yet with inconsistent reports of the direction of BOLD changes, implying a dependence on experimental and methodological factors. In mice, OT receptor-expressing neurons in the medial amygdala have been found to mediate olfactory-based social familiarity effects (*26*). Here, though, stranger touch activated the amygdala in a general fashion (Fig 3C; Table S5), supporting the proposal that its prominent role in human and nonhuman primate OT studies may be indirect (*93, 94*). The covariant relationship of ITG with hedonic and social evaluation ratings and amygdala activation, especially for partner touch in the second encounter (Fig 3D), suggests a potential role for this region in maintaining receptivity to touch following contextual shifts.

In recent decades, a specific subtype of C afferent nerves (C-tactile or CT) has become increasingly implicated in affective touch (*52*,*54*). CT afferent signaling has been proposed to interact with OT modulation (*5*). However, we found no supporting evidence for such a relationship; neither somatosensory cortices nor the posterior insular/inferior parietal regions selectively associated with CT stimulation on the arm vs palm skin covaried with OT changes, who delivered the touch, or in what order he was presented to the participant. Instead, somatosensory and posterior insular cortices showed general responses to touch in a conjunction across all touch conditions (Table S6). IN-OT has been observed to increase the pleasantness of touch (*8*), and individuals with higher salivary OT levels have reported greater touch pleasantness (*95*). Neurons in deep (II-X), but not superficial (I), layers of the spinal cord project to the caudo-dorsal part of the PVN (*96*), where parvocellular OT neurons are located, and non-peptidergic C afferent neurons have been observed to express the OT receptor (OXTR; *96*). Yet this work has so far not identified OXTR expression in dorsal horn neurons of the spinothalamic tract projecting to the specific thalamic pathways putatively shared by CT afferents (*97*, *98*). Further research is therefore needed to dissect the relevant spinothalamic sensory circuitry and its relation to touch-driven OT effects.

The measurement of peripheral OT in humans comes with caveats, as does its relationship with central mechanisms of release. Different methods for detecting plasma OT have yielded different and sometimes uncorrelated sets of value ranges, with measurement issues centering around the selectivity with which testing components detect bound or unbound protein, or whole or fragmentary molecules. Cerebrospinal fluid (CSF) has consistently been found to correlate more strongly with brain OT levels than does plasma OT, while CSF and plasma OT measurements have shown weak or no correlation. However, at least a proportion of inconsistencies in reported findings may depend on the investigation of basal levels, in the absence of acute stimuli more likely to evoke coordinated, biologically-meaningful responses across the central and peripheral nervous systems, which may also include bioactivity of fragments (*100*). In this study, within-subject serial sampling allowed us to assess evoked OT changes with respect to individual baselines. The covariation of these changes and its temporal patterns with BOLD points to a relationship between the central and peripheral effects of ecologically-evoked endogenous OT, but cannot directly demonstrate it.

### Conclusions

These findings offer a methodological and conceptual bridge between stimulus-driven and context-sensitive frameworks of endogenous OT modulation of the brain during social interactions. OT’s more general role in social interactions may involve reinforcing the *status quo* with respect to familiar individuals, yet also selectively modulating brain-body responses during interactions with unfamiliar individuals. Such selective differences are likely modulable across subsequent encounters, and may thus calibrate neural and behavioral receptivity in social interactions over time, whether mediated by touch or another channel such as vision or speech. Network hubs in parietotemporal pathways, alongside precuneus and retrosplenial cortex, may mediate such influence, turning the “dimmer” of neural processing up or down depending on past and current social context.

## Supporting information

Handlin Novembre et al Supplementary

## Acknowledgments

The authors thank Åsa Axén and Gisela Öhnström for blood sample collection, Kerstin Uvnäs-Moberg, Maria Petersson, Stephanie Preston, and Ellen Lumpkin for valuable discussion, and Paul Hamilton and Irene Perini for assistance with AFNI software.

## Author contributions

I.M. and L.H. designed the study; G.N., L.H., H.L. and I.M collected the data; G.N., L.H., H.L., and R.K. analyzed the data; I.M., G.N., and L.H. wrote the paper.

## Data and materials availability

Anonymized fMRI data are available upon request; blood hormone data availability is limited by ethical, data protection, and materials transfer regulations.

## References

1. Tang, Y., et al. (2020). Social touch promotes interfemale communication via activation of parvocellular oxytocin neurons. Nat Neurosci, 23: 1125–1137.

2. Kurosawa, M., et al. (1995). Massage-like stroking of the abdomen lowers blood pressure in anesthetized rats: influence of oxytocin. J Auton Nerv Syst, 56: 26–30.

3. Feldman, R. (2012). Oxytocin and social affiliation in humans. Horm Behav 61: 380–391.

4. Uvnäs-Moberg, K., L. Handlin and M. Petersson (2015). Self-soothing behaviors with particular reference to oxytocin release induced by non-noxious sensory stimulation. Frontiers in Psychology 5: 1–16.

5. Walker, S.C. et al. (2017). C-tactile afferents: Cutaneous mediators of oxytocin release during affiliative tactile interactions? Neuropeptides 64, 27–38.

6. Li, Q. et al. (2019). Foot massage evokes oxytocin release and activation of orbitofrontal cortex and superior temporal sulcus. Psychoneuroendocrinol 101, 193–203.

7. Kreuder, A.K. et al. (2017). How the brain codes intimacy: The neurobiological substrates of romantic touch. Hum Brain Mapp, 38, 4525–4534.

8. Chen, Y., et al (2020). Oxytocin increases the pleasantness of affective touch and orbitofrontal cortex activity independent of valence. Eur Neuropsychopharmacol, doi:10.1016/j.euroneuro.2020.08.003

9. M. Mitre et al. (2018). Oxytocin modulation of neural circuits. Curr Top Beh Neurosci 35, 31.53

10. Knobloch, H. S. et al., (2012) Evoked axonal oxytocin release in the central amygdala attenuates fear response. Neuron 73: 553–566.

11. Oettl, L.L. et al. (2016) Oxytocin enhances social recognition by modulating cortical control of early olfactory processing. Neuron 90: 609–621.

12. Burbach, J.P.H. et al (2006). Oxytocin: Synthesis, Secretion and Reproductive Functions. In: Knobil and Neill’s Physiology of Reproduction. 3 ed: Elsevier).

13. Wang P, Wang SC, Liu X, Jia S, Wang X, Li T, Yu J, Parpura V, Wang YF. (2022). Neural Functions of Hypothalamic Oxytocin and its Regulation. ASN Neuro:17590914221100706.

14. Yu H, Miao W, Ji E, Huang S, Jin S, Zhu X, Liu MZ, Sun YG, Xu F, Yu X. (2022). Social touch-like tactile stimulation activates a tachykinin 1-oxytocin pathway to promote social interactions. Neuron 110(6):1051–1067.e7.

15. Qin, J. et al., (2009). Oxytocin receptor expressed on the smooth muscle mediates the excitatory effect of oxytocin on gastric motility in rats. Neurogastroenterol. Motil. 21, 430–438.

16. Filippi, S. et al. (2003). Role of oxytocin in the ejaculatory process. J. Endocrinol. Invest. 26, 82–86.

17. Uvnas-Moberg, K. et al. (1998). Oxytocin may mediate the benefits of positive social interaction and emotions. Psychoneuroendocrinol 23, 819–35.

18. Walum H, Young LJ. (2018). The neural mechanisms and circuitry of the pair bond. Nat Rev Neurosci 19:643–654.

19. Algoe, S.B. et al (2017). Oxytocin and social Bonds: the role of oxytocin in perceptions of romantic partners’ bonding behavior. Psychol Sci, 28:1763–1772.

20. T. Rehn et al. (2014). Dogs’ endocrine and behavioural responses at reunion are affected by how the human initiates contact. Physiol Behav, 124, 45–53.

21. Nagasawa, M. et al. (2015). Social evolution. Oxytocin-gaze positive loop and the coevolution of human-dog bonds, Science 348: 333–336.

22. O. Dal Monte, et al (2017). Oxytocin under opioid antagonism leads to supralinear enhancement of social attention. PNAS 114, 5247–5252.

23. Ferretti, V. et al (2019). Oxytocin signaling in the central amygdala modulates emotion discrimination in mice. Curr Biol 17:1938–1953.e6.

24. Ferguson, J.N. et al. (2001). Oxytocin in the medial amygdala is essential for social recognition in the mouse. J. Neurosci. 21, 8278–8285.

25. Quintana D.S., (2020). An allostatic theory of oxytocin. Trends Cogn. Sci. 24: 515–528.

26. J.A. Bartz, et al. (2011). Social effects of oxytocin in humans: context and person matter. Trends Cogn Sci 15: 301–309.

27. J.P. Burkett et al. (2016). Oxytocin-dependent consolation behavior in rodents. Science 351: 375–378

28. Rickenbacher, E et al (2017). Freezing suppression by oxytocin in central amygdala allows alternate defensive behaviours and mother-pup interactions. Elife 6, doi:10.7554/eLife.24080

29. LoParo, D. et al (2016). Rigorous tests of gene-environment interactions in a lab study of the oxytocin receptor gene (OXTR), alcohol exposure, and aggression. Am J Med Genet B Neuropsychiatr Genet, 171: 589–602.

30. Hovey, D. et al (2016). Antisocial behavior and polymorphisms in the oxytocin receptor gene: findings in two independent samples. Mol Psychiatry 21: 983–988.

31. Anpilov S. et. al. (2020). Wireless Optogenetic Stimulation of Oxytocin Neurons in a Semi-natural Setup Dynamically Elevates Both Pro-social and Agonistic Behaviors. Neuron. 107: 644–655.

32. Huang H. et al (2014). Chronic and acute intranasal oxytocin produce divergent social effects in mice. Neuropsychopharmacology 39: 1102–1114.

33. Leng, G. and Russell, J.A. (2019). The osmoresponsiveness of oxytocin and vasopressin neurones: Mechanisms, allostasis and evolution. J. Neuroendocrinol. 31, e12662.

34. Anacker, A.M. and Beery, A.K. (2013). Life in groups: the roles of oxytocin in mammalian sociality. Front Behav Neurosci 7:185.

35. De Dreu C.K., et. al. (2010). The neuropeptide oxytocin regulates parochial altruism in intergroup conflict among humans. Science 328: 1408–1411.

36. Heinrichs, M. et al (2003). Social support and oxytocin interact to suppress cortisol and subjective responses to psychosocial stress. Biol Psychiatry 54:1389–98.

37. Di Simplicio, J. (2009). Oxytocin enhances processing of positive versus negative emotional information in healthy male volunteers. Psychopharmacol 23:241–8.

38. Fischer-Shofty, M., et. al. (2010). The effect of intranasal administration of oxytocin on fear recognition. Neuropsychologia 48: 179–184.

39. Petrovic, P. et al (2008). Oxytocin attenuates affective evaluations of conditioned faces and amygdala activity. J Neurosci 28: 6607–6615.

40. Rimmele, U. et al (2009). Oxytocin makes a face in memory familiar. J Neurosci 29: 38–42.

41. Domes, G., et. al. (2007). Oxytocin improves “mind-reading” in humans. Biol Psychiatry 61: 731–733.

42. Ditzen, B., et. al. (2009). Intranasal oxytocin increases positive communication and reduces cortisol levels during couple conflict. Biol Psychiatry, 65(9), 728–731.

43. Scheele, D. et. al. (2012). Oxytocin modulates social distance between males and females. J Neurosci 32: 16074–16079.

44. Scheele, D., et. al. (2013). Oxytocin enhances brain reward system responses in men viewing the face of their female partner. Proc Natl Acad Sci U S A 110: 20308–20313.

45. A.K. Kreuder et al., How the brain codes intimacy: The neurobiological substrates of romantic touch. Hum Brain Mapp, 38, 4525–4534 (2017).

46. Kreuder, A. K. et al (2019). Oxytocin enhances the pain-relieving effects of social support in romantic couples. Hum Brain Mapp, 40: 242–251.

47. D. Wang, et al. (2017). Neural substrates underlying the effects of oxytocin: a quantitative meta-analysis of pharmaco-imaging studies. Soc Cogn Affect Neurosci 12: 1565–73.

48. C.F. Zink & A. Meyer-Lindenberg, A. (2012). Human neuroimaging of oxytocin and vasopressin in social cognition. Horm Behav 61, 400–9.

49. Striepens, N., et. al. (2013). Elevated cerebrospinal fluid and blood concentrations of oxytocin following its intranasal administration in humans. Sci Rep 3: 3440.

50. Churchland, P. S., & Winkielman, P. (2012). Modulating social behavior with oxytocin: how does it work? What does it mean? Horm Behav 61: 392–399.

51. McCullough, M. E. et al (2013). Problems with measuring peripheral oxytocin: can the data on oxytocin and human behavior be trusted? Neurosci Biobehav Reviews 37: 1485–1492.

52. Löken, L. S., et. al. (2009). Coding of pleasant touch by unmyelinated afferents in humans. Nat Neurosci 12: 547–548.

53. Morrison, I. et al (2010). The skin as a social organ. Experimental Brain Research 204: 305–314.

54. Olausson, H. et. al. (2002). Unmyelinated tactile afferents signal touch and project to insular cortex. Nat Neurosci 5: 900–904.

55. Shansky, R.M., Murphy, A.Z. (2021). Considering sex as a biological variable will require a global shift in science culture. Nat Neurosci 24:457–464.

56. S. Filippi et al (2003). Role of oxytocin in the ejaculatory process. J Endocrinol Invest 26(3 Suppl):82–6.

57. A. Salonia (2005). Menstrual cycle-related changes in plasma oxytocin are relevant to normal sexual function in healthy women. Horm Behav 47:164-9. 2005 Feb;47(2):164–9.

58. J.B. Becker et el. (2016). Female rats are not more variable than male rats: a meta-analysis of neuroscience studies. Biol Sex Differences 7:34.

59. Prendergast BJ, Onishi KG, Zucker I. Female mice liberated for inclusion in neuroscience and biomedical research. Neurosci Biobehav Rev. 2014;40:1–5.

60. Itoh Y, Arnold AP. Are females more variable than males in gene expression? Meta-analysis of microarray datasets. Biol Sex Differ. 2015;6:18.

61. J.L. Funk and R.D. Rogge, Testing the ruler with item response theory: Increasing precision of measurement for relationship satisfaction with the Couples Satisfaction Index. Journal of Family Psychology 21, 572–583 (2007).

62. C.D: Spielberger et al., Manual for the State-Trait Anxiety Inventory. Palo Alto, CA: Consulting Psychologists Press (1983).

63. Cox, R. W., Chen, G., Glen, D. R., Reynolds, R. C. & Taylor, P. A. fMRI clustering and false-positive rates. PNAS 114, E3370–e3371 (2017).

64. T. A. Myers, Goodbye, listwise deletion: presenting hot deck imputation as an easy and effective tool for handling missing data, communication methods and measures, 5, 297–310 (2011).

65. LoParo, D. et al (2016). Rigorous tests of gene-environment interactions in a lab study of the oxytocin receptor gene (OXTR), alcohol exposure, and aggression. Am J Med Genet B Neuropsychiatr Genet, 171: 589–602.

66. Scheele, D. et. al. (2012). Oxytocin modulates social distance between males and females. J Neurosci 32: 16074–16079.

67. Scheele, D., et. al. (2013). Oxytocin enhances brain reward system responses in men viewing the face of their female partner. Proc Natl Acad Sci U S A 110: 20308–20313.

68. A.K. Kreuder et al., How the brain codes intimacy: The neurobiological substrates of romantic touch. Hum Brain Mapp, 38, 4525–4534 (2017).

69. Kreuder, A. K. et al (2019). Oxytocin enhances the pain-relieving effects of social support in romantic couples. Hum Brain Mapp, 40: 242–251.

70. Lancaster K, Carter CS, Pournajafi-Nazarloo H, Karaoli T, Lillard TS, Jack A, Davis JM, Morris JP, Connelly JJ. (2015). Plasma oxytocin explains individual differences in neural substrates of social perception. Front Hum Neurosci 9:132.

71. D. Wang, et al. (2017). Neural substrates underlying the effects of oxytocin: a quantitative meta-analysis of pharmaco-imaging studies. Soc Cogn Affect Neurosci 12: 1565–73.

72. C.F. Zink & A. Meyer-Lindenberg, A. (2012). Human neuroimaging of oxytocin and vasopressin in social cognition. Horm Behav 61, 400–9.

73. Shamay-Tsoory S.G., and Abu-Akel A. (2016). The Social Salience Hypothesis of Oxytocin. Biol. Psychiatry 79: 194–202.

74. Z.V. Johnson et al. (2017). Oxytocin receptors modulate a social salience neural network in male prairie voles. Horm Behav 87: 16–24.

75. Sripada, C. S., et. al. (2013). Oxytocin enhances resting-state connectivity between amygdala and medial frontal cortex. Int J Neuropsychopharmacol 16: 255–260.

76. Martins, D. A. et al (2020). Effects of route of administration on oxytocin-induced changes in regional cerebral blood flow in humans. Nat Commun 11: 1160.

77. Valstad, M. et al (2017). The correlation between central and peripheral oxytocin concentrations: A systematic review and meta-analysis. Neurosci Biobeh Rev 78: 117–124.

78. Pow, D.V. and Morris, J.F. (1989). Dendrites of hypothalamic magnocellular neurons release neurohypophysial peptides by exocytosis. Neuroscience 32: 435–439.

79. D.S. Margulies et al (2009). Precuneus shares intrinsic functional architecture in humans and monkeys. PNAS 106: 20069–74.

80. J. Kumar et al. (2019). Oxytocin modulates the effective connectivity between the precuneus and the dorsolateral prefrontal cortex. Eur Arch Psychiatry Clin Neurosci: s00406-019-00989.

81. Petersson, M., & Uvnas-Moberg, K. (2003). Systemic oxytocin treatment modulates glucocorticoid and mineralocorticoid receptor mRNA in the rat hippocampus. Neurosci Lett 343: 97–100.

82. Vargas-Martinez, F., et. al. (2014). Neuropeptides as neuroprotective agents: Oxytocin a forefront developmental player in the mammalian brain. Prog Neurobiol 123: 37–78.

83. J. Winter & B. Jurek (2019). The interplay between oxytocin and the CRF system: regulation of the stress response. Cell Tissue Res 375: 85–91.

84. Grewen K.M. et al (2005). Effects of partner support on resting oxytocin, cortisol, norepinephrine, and blood pressure before and after warm partner contact. Psychosom Med 67:531–8.

85. Ditzen, B. et al (2009). Intranasal oxytocin increases positive communication and reduces cortisol levels during couple conflict. Biol Psychiatry 65: 728–731.

86. Davidovic, M., et. al. (2016). Posterior superior temporal sulcus responses predict perceived pleasantness of skin stroking. Frontiers in Human Neuroscience 10: 432.

87. Kaiser, M. D., et. al. (2016). Brain mechanisms for processing affective (and nonaffective) touch are atypical in autism. Cereb Cortex 26: 2705–2714.

88. Voos, A. C., et. al. (2013). Autistic traits are associated with diminished neural response to affective touch. SCAN 8: 378–386.

89. Liu, Y. et al. (2019). Oxytocin modulates social value representations in the amygdala. Nat Neurosci 22: 633–641

90. Motoki, K., et al., (2016). Are plasma oxytocin and vasopressin levels reflective of amygdala activation during the processing of negative emotions? A preliminary study. Front Psychol 7: 480.

91. Sripada, C.S. et al. (2013). Oxytocin enhances resting-state connectivity between amygdala and medial frontal cortex. Int J Neuropsychopharmacol 16, 255–260.

92. Kirsch, P., et. al., Oxytocin modulates neural circuitry for social cognition and fear in humans. J Neurosci, 25(49), 11489–11493 (2005).

93. P.T. Putnam et al. (2018). Bridging the gap between rodents and humans: The role of non-human primates in oxytocin research. Am J Primatol 80: e22756.

94. M. Eckstein et al. (2017). Oxytocin differentially alters resting state functional connectivity between amygdala subregions and emotional control networks: Inverse correlation with depressive traits. NeuroImage 149: 458–67.

95. Portnova, G.V. et al. (2020). Perceived pleasantness of gentle touch in healthy individuals is related to salivary oxytocin response and EEG markers of arousal. Experimental Brain Research

96. Gauriau, C., & Bernard, J. F. (2004). A comparative reappraisal of projections from the superficial laminae of the dorsal horn in the rat: the forebrain. J Comp Neurol 468: 24–56.

97. Nersesyan, Y. et al (2017). Oxytocin Modulates Nociception as an Agonist of Pain-Sensing TRPV1. Cell Rep 21: 1681–1691.

98. Moreno-Lopez, Y. et al (2013). Identification of oxytocin receptor in the dorsal horn and nociceptive dorsal root ganglion neurons. Neuropeptides 47: 117–123.

99. Andrew, D. (2010). Quantitative characterization of low-threshold mechanoreceptor inputs to lamina I spinoparabrachial neurons in the rat. J Physiol 588: 117–124.

100. Uvnäs-Moberg, K. et. At. (2019). Oxytocin is a principal hormone that exerts part of its effects by active fragments. Med Hypotheses 133: 109394.S

