## Supplementary material for "Human endogenous oxytocin and its neural correlates show adaptive responses to social touch based on recent social context": Handlin Novembre et al Supplementary

#### Supplementary Methods

*Piloting and sample size selection.* Since this experiment involved several different types of measures (OT, BOLD, and behavioral ratings), we selected our sample size with the aim of balancing OT and BOLD, while including a margin for potential data loss in serial plasma sampling. The overall sample size of 30+ was therefore chosen on the basis of multiple considerations. Analysis of pilot data indicated that a sample size of n = 80 (40 per group) is more than sufficient to detect group OT differences between partner-first and stranger-first presentation order at 80% power, with alpha = 0.05. Further, a minimum sample size of n = 24 has been estimated as sufficient to detect BOLD effects in a block design at 80% power, with alpha = 0.002 (*1*). Data analysis did not occur until all data was collected and had roughly equal counterbalancing among participants with complete hormone samples (final OT sample n = 27; final BOLD sample for general linear model n = 35; see Table S1).

*OT regressor*

To discover the neural correlates of endogenous OT modulation regardless of toucher order, we used each participant's plasma OT levels, for each sample collected during the functional runs, to create a regressor that tested for activation corresponding with the overall temporal pattern of the endogenous OT response. N=23 participants had complete data series for both functional runs (see s1). All missing data-points (points between samples) were obtained through linear interpolation and the resulting function was convolved with the canonical HRF. By adding time lags of 1, 1.5, 2, 2.5 and 3 min, this regressor allowed identification of activation preceding the plasma OT samples in these five time windows. Thus, it captured potential central modulation corresponding to the peripheral changes in plasma OT observed after various delays.

This analysis revealed brain areas showing significant interaction between order and stroker at both 2 and 2.5 minutes, as a result of higher activity during stranger first than partner first, but no difference between stranger second and partner second. No activations were observed for the remaining time lags.

*2 min.* Two minutes before measured changes in oxytocin levels, this pattern of activity was found in bilateral precuneus, right posterior cingulate cortex, right postcentral gyrus, right parietal operculum, right SMG, and left inferior parietal lobule (p = 0.002; Table s4).

*2.5 min.* Two and half minutes before measured changes in oxytocin levels, the same pattern of activity was observed in bilateral precuneus and right paracentral lobule (all at p = 0.002, Table s4).

#### Results

##### Oxytocin-independent analyses

*Conjunctions*

*Initial encounter: Partner first ∩ stranger first:* Brain areas showing increased activity in the first run, regardless of stroker identity, were: right PO, bilateral TP, right parahippocampal gyrus, right inferior parietal lobule (IPL), and bilateral cerebellum. There were also common deactivations in clusters located in the right postcentral and precentral gyri.

*Second encounter: Partner second ∩ stranger second:* Brain areas showing increased activity during the second run, regardless of stroker identity, were: right cerebellum, left caudate and left putamen. There was also a common deactivation in the right precentral gyrus.

*Partner first: Partner first ∩ stranger second:* Specific activation increases within the partner first group fell in right precentral gyrus, right SPL, right superior temporal gyrus (STG), right superior frontal gyrus, bilateral cerebellum, and bilateral middle temporal gyrus (MTG). There were also common deactivations in clusters located in the right postcentral and precentral gyri.

*Stranger first: Stranger first ∩ partner second:* Specific activation increases within the stranger first group fell in right inferior parietal lobule (IPL), right cerebellum, left caudate, bilateral MCC. There was also a common deactivation in the right precentral gyrus.

*Partner: Partner first ∩ partner second:* Brain areas showing specific activation increases during partner touch, independently of the group, were: left postcentral gyrus, and right cerebellum. There was also a common deactivation in the right precentral gyrus.

*Stranger: Stranger first ∩ stranger second:* Brain areas showing specific activation increases during stranger touch, independently of the group, were: right precentral gyrus, right hippocampus, right putamen, bilateral cerebellum, left caudate, and TP. There was also a common deactivation in the right postcentral gyrus.

#### Supplementary references

1. Desmond JE & Glover GH (2002). Estimating sample size in functional MRI (fMRI) neuroimaging studies: statistical power analyses. *J Neurosci Methods* 118: 115-128.

#### Supplementary Tables:

**Table S1**. Participants included in each analysis based on analyzable data. OT = Oxytocin, CORT = Cortisol, LME = Linear Mixed Effects Model (neuroimaging), Regr. = Regressor (neuroimaging).

| Group | Participant No. | Analyzable samples | | Analyses | | | | | | | | |
|---|---|---|---|---|---|---|---|---|---|---|---|---|
| | | OT | CORT | Plasma | | LME | | T test Covariate | | OT Regr. | ANOVA | Conjunctions |
| | | | | OT | CORT | OT | CORT | OT | CORT | | | |
| Partner First - Stranger Second | 1 | 1,2,3,4 | 1,2,4 | - | - | - | - | - | - | - | - | P |
| | 2 | 1,2,3,4,5,6,7,8,9 | 1,2,4,5,6,8,9 | X | X | X | X | X | X | X | X | X |
| | 3 | 1,2,3,4,5 | 1,2,4,5 | - | - | - | - | - | - | - | - | P |
| | 4 | 1,2,3,4,5,6,7,8,9 | 1,2,4,5,6,8,9 | X | X | X | X | X | X | X | X | X |
| | 5 | 1,2,3,4,5,6,7,8,9 | 1,2,4,5,6,8,9 | X | X | X | X | X | X | X | X | X |
| | 6 | 1,2,3,4,5,7 | 1,2,4,5,6,8 | X | X | X | X | X | X | - | X | X |
| | 7 | 1,2,3,4,5,6,7,8,9 | 1,2,4,5,6,8,9 | X | X | X | X | X | X | X | X | X |
| | 8 | - | - | - | - | - | - | - | - | - | X | X |
| | 9 | 1,2,3,4,5,6,7,8,9 | 1,2,4,5,6,8,9 | X | X | X | X | X | X | X | X | X |
| | 10 | - | - | - | - | - | - | - | - | - | X | X |
| | 11 | 1,2,3,4,5,6,7,8,9 | 1,2,4,5,6,8,9 | X | X | X | X | X | X | X | X | X |
| | 12 | 1 | 1 | - | - | - | - | - | - | - | X | X |
| | 13 | - | - | - | - | - | - | - | - | - | X | X |
| | 14 | 1,3,4,5,6,9 | 1,2,4,5,6,9 | - | - | - | - | - | - | - | - | - |
| | 15 | 1,2,3,4,5,6,7,8,9 | 1,2,4,5,6,8,9 | X | X | X | X | X | X | X | X | X |
| | 16 | - | - | - | - | - | - | - | - | - | - | - |
| | 17 | 1,2,3,4,5,6,7,8,9 | 1,2,4,5,6,8,9 | X | X | X | X | X | X | X | X | X |
| | 18 | 1,2,3,4,5,6,7,8,9 | 1,2,4,5,6,8,9 | X | X | X | X | X | X | X | X | X |
| | 19 | 1,2,3,4,5,6,7,8,9 | 1,2,4,5,6,8,9 | X | X | X | X | X | X | X | X | X |
| | 20 | 1,2,3,4,5,6,7,8,9 | 1,2,4,5,6,8,9 | X | X | X | X | X | X | X | X | X |
| | 21 | 1,4,5 | 1,4,5 | - | - | - | - | - | - | - | - | - |
| | 22 | 1,2,3,4,5,6,7,8,9 | 1,2,4,5,6,8,9 | X | X | X | X | X | X | X | X | X |
| | 23 | 1,2,3,4,5,6,7,8 | 1,2,4,5,6,8,9 | X | X | X | X | X* | X | X | X | X |
| Stranger First - Partner Second | 1 | 1,2,3,4,5,6,7,8,9 | 1,2,4,5,6,8,9 | X | X | X | X | X* | X | X | X | X |
| | 2 | 1,2,3,4,5,6,7,8,9 | 1,2,4,5,6,8,9 | X | X | X | X | X | X | X | X | X |
| | 3 | 1,2,3,4,5,6,7,8,9 | 1,2,4,5,6,8,9 | X | X | X | X | X | X | X | X | X |
| | 4 | - | 1,2 | - | - | - | - | - | - | - | X | X |
| | 5 | 1,2,3,4,5,6,7,8,9 | 1,2,4,5,6,8,9 | X | X | X | X | X | X | X | X | X |
| | 6 | - | - | - | - | - | - | - | - | - | - | - |
| | 7 | 2,3,4,5,6,7,8,9 | 2,4,5,6,8,9 | - | - | - | - | - | - | - | X | X |
| | 8 | 1 | - | - | - | - | - | - | - | - | X | X |
| | 9 | 1,2,3,4,5,6,8,9 | 1,2,4,5,6,8,9 | X | X | X | X | X | X | - | X | X |
| | 10 | 1,2,3,4,5,7,8,9 | 1,2,4,5,6,8,9 | X | X | X | X | X | X | - | X | X |
| | 11 | 1,2,3,4,5,6,7,8,9 | 1,2,4,5,6,8,9 | X | X | X | X | X | X | X | X | X |
| | 12 | 1,2,3,4,5,6,7,8 | 1,2,4,5,6,8,9 | X | X | X | X | X | X | X | X | X |
| | 13 | - | - | - | - | - | - | - | - | - | X | X |
| | 14 | 1,2,3,4,5,6,7,8,9 | - | X | - | X | - | X | - | X | X | X |
| | 15 | 1,2,3,4,5,6,7,8,9 | 1,2,4,5,6,8,9 | X | X | X | X | X | X | X | X | X |
| | 16 | 3 | 1,2,4 | - | - | - | - | - | - | - | - | - |
| | 17 | 1,2,3,4,5,6,7,8 | 1,2,4,5,6,8,9 | X | X | X | X | X | X | X | X | X |
| | 18 | 1,2,3,5,7,8 | 1,2,4,5,6,8,9 | X | X | X | X | X | X | - | X | X |
| | 19 | 1,2,3,4,5,6,7,8,9 | 1,2,4,5,6,8,9 | X | X | X | X | X | X | X | X | X |

* = Participants excluded from the from Hypothalamus ROI analysis for signal dropout in that region.

P = only the Partner run is included in this analysis.

S = only the Stranger run is included in this analysis.

**Table S2.** Linear mixed-effects modeling with factors toucher (partner, stranger), order (first or second encounter), and peak OT changes. All contrasts thresholded at P<0.002, cluster-size thresholded at alpha=0.05 FWE for n = 35 complete functional datasets. For each cluster under each contrast heading, size, location, maximum F, and MNI coordinates (x, y, z) are given.

| Cluster number (size) | Cluster location | F (x, y, z) |
|---|---|---|
| ***Interaction: Toucher x Order x Oxytocin*** | | |
| Cluster #1 (91) | Right Superior Occipital Gyrus | 29.77 (22, -98, 19) |
| | Right Cuneus | 20.10 (16, -101, 10) |
| ***Interaction: Toucher x Oxytocin*** | | |
| Cluster #1 (212) | Left Anterior Cingulate Cortex | 23.42 (-11, 49, 1) |
| | Right Mid Orbital Gyrus | 23.17 (4, 64, -2) |
| | Right Mid Orbital Gyrus | 22.91 (13, 49, -5) |
| | Right Anterior Cingulate Cortex | 20.56 (10, 49, 13) |
| Cluster #2 (150) | Right Supramarginal Gyrus | 24.73 (49, -53, 34) |
| Cluster #3 (144) | Right Middle Temporal Gyrus | 28.44 (46, -2, -26) |
| | Right Temporal Pole | 26.32(40, 19, -32) |
| | Right Medial Temporal Pole | 21.63 (49, 10, -32) |
| Cluster #4 (75) | Right Superior Frontal Gyrus | 33.76 (22, 25, 61) |
| | Right Superior Frontal Gyrus | 27.70 (22, 16, 67) |
| | Right Superior Medial Gyrus | 22.49 (13, 31, 58) |

***Interaction: Toucher x Oxytocin***

| | | |
|---|---|---|
| Cluster #1 (88) | Right Inferior Temporal Gyrus | 22.76 (46, -5, -32) |
| | Right Temporal Pole | 20.42 (40, 19, -32) |

**Table S3.** Paired T-tests for partner vs stranger during each of 2 functional runs (first, second), modeled with linear mixed effects and weighted by individual peak OT levels as covariate. All contrasts thresholded at P<0.002, cluster-size thresholded at alpha=0.05 FWE for n = 35 complete functional datasets. For each cluster under each contrast heading, size, location, maximum T, and MNI coordinates (x, y, z) are given. * = region of interest analysis.

| Cluster number (size) | Peak location | T (x, y, z) |
|---|---|---|
| **Partner First > Stranger First** | | |
| Cluster #1 (72) | Left Raphe | 5.76 (-11, -47, -41) |
| | Right Raphe | 4.46 (7, -44, -38) |
| Cluster #2 (15) | Left Hypothalamus* | 4.16 (-2, -2, -8) |
| **Partner Second > Stranger Second** | | |
| Cluster #1 (223) | Right Angular Gyrus | 5.09 (40, -53, 25) |
| | Right Angular Gyrus | 4.70 (52, -62, 31) |
| | Right Supramarginal Gyrus | 4.39 (46, -44, 28) |
| | Right Superior Temporal Gyrus | 4.02 (52, -44, 16) |
| Cluster #2 (166) | Right Middle Temporal Gyrus | 5.30(46, 4, -29) |
| | Right Medial Temporal Pole | 5.09 (43, 16, -35) |
| | Right Middle Temporal Gyrus | 4.77 (55, -2, -29) |
| Cluster #2 (163) | Right Anterior Cingulate Cortex | 4.74(13, 49, 10) |
| | Right Superior Medial Gyrus | 4.61 (7, 67, 1) |

| | | |
|---|---|---|
| | Right Mid Orbital Gyrus | 4.14(10, 49, -5) |
| | Left Mid Orbital Gyrus | 4.04(-8, 55, -2) |
| Cluster #4 (88) | Right Superior Frontal Gyrus | 6.07 (25, 19, 64) |
| | Right Superior Frontal Gyrus | 5.66 (19, 28, 61) |

**Table S4.** Regressor created by linear interpolation of serial OT samples, convolved with canonical HRF and modeled with factors toucher (partner, stranger) and order (first or second encounter), at time points 2 and 2.5 min preceding plasma sample collection. All contrasts thresholded at P<0.002, cluster-size thresholded at alpha=0.05 FWE for N = 24 complete functional datasets. For each cluster under each contrast heading, size, location, maximum F, and MNI coordinates (x, y, z) are given.

### 2 minutes preceding sample collection

| Cluster number (size) | Peak location | F (x, y, z) |
|---|---|---|
| ***Interaction: Toucher x Order*** | | |
| Cluster #1 (250) | Left Precuneus | 33.04 (-8, -62, 67) |
| | Right Precuneus | 25.65 (10, -65, 58) |
| | Right Precuneus | 20.84 (16, -44, 49) |
| | Right Precuneus | 20.70 (13, -50, 58) |
| | Left Precuneus | 17.03 (-14, -50, 79) |
| Cluster #2 (87) | Right Postcentral Cingulate Cortex | 25.91 (10, -53, 10) |
| | Left Precuneus | 24.83 (-5, -53, 13) |
| Cluster #3 (63) | Left Inferior Parietal Lobule | 37.21 (-50, -54, 63) |

### 2.5 minutes preceding sample collection

***Interaction: Toucher x Order***

| | | |
|---|---|---|
| Cluster #1 (234) | Right Precuneus | 27.24 (1, -47, 61) |

| | | |
|---|---|---|
| | Right Precuneus | 26.64 (16, -44, 52) |
| | Left Precuneus | 21.65 (-2, -65, 55) |
| | Left Precuneus | 20.62 (-2, -80, 46) |
| | Left Precuneus | 19.87 (-8, -59, 67) |
| | Left Precuneus | 16.31 (-8, -47, 55) |
| | Right Paracentral Lobule | 14.71 (7, -35, 52) |

**Table S5.** Conjunction analyses showing common activations for partner and order factors (partner, stranger, first, second). All contrasts thresholded at P<0.002, cluster-size thresholded at alpha=0.05 FWE for n = 35 complete functional datasets. For each cluster under each contrast heading, size, location, maximum T, and MNI coordinates (x, y, z) are given. Negative BOLD in boldface.

***Partner First ∩ Stranger Second ∩ Stranger First ∩ Partner Second***

| Cluster number (size) | Peak location | T (x, y, z) |
|---|---|---|
| Cluster #1 (2264) | Left Supramarginal Gyrus | 11.68 (-56, -23, 43) |
| | Left Postcentral Gyrus | 11.18 (-50, -32, 52) |
| | Left Postcentral Gyrus | 10.80 (-35, -41, 61) |
| | Left Precentral Gyrus | 10.79 (-29, -14, 67) |
| | Left Postcentral Gyrus | 10.48 (-41, -35, 67) |
| | Left Supramarginal Gyrus | 10.47 (-50, -32, 25) |
| | Left Superior Parietal Lobule | 8.99 (-23, -44, 73) |
| | Left Parietal Operculum | 8.56 (-44, -5, 7) |
| | Left Supplementary Motor Area | 7.56 (-5, -5, 58) |
| | Left Precentral Gyrus | 7.12 (-59, 7, 31) |
| Cluster #2 (356) | Right Inferior Frontal Gyrus | 6.16 (58, 13, 28) |
| | Right Insula | 5.74 (43, 16, -5) |
| | Right Inferior Frontal Gyrus | 5.64 (58, 16, 7) |
| | Right Inferior Frontal Gyrus | 5.49 (40, 31, 1) |
| | Right Insula | 5.36 (43, 1, 1) |
| Cluster #3 (332) | Right Supramarginal Gyrus | 6.74 (58, -20, 22) |
| | Right Supramarginal Gyrus | 5.90 (52, -26, 34) |
| | Right Supramarginal Gyrus | 5.75 (37, -35, 43) |
| Cluster #4 (118) | Right Cerebellum | 8.82 (25, -50, -26) |

| | | |
|---|---|---|
| | Right Cerebellum | 5.07 (10, -59, -21) |
| Cluster #5 (115) | Right Cerebellum | 8.47 (19, -62, -53) |
| Cluster #6 (83) | Right Precentral Gyrus | **-5.96** (31, -26, 61) |
| | Right Precentral Gyrus | **-5.48** (37, -20, 52) |
| Cluster #7 (78) | Right Supplementary Motor Area | 6.95 (4, 7, 55) |
| Cluster #8 (14) | Left Insula | 4.50 (-29, 10, -20) |
| Cluster #9 (11) | Left Middle Cingulate Cortex | 5.34 (-8, 13, 43) |
| Cluster #10 (9) | Right Amygdala | 4.71 (31, 4, -20) |
| Cluster #11 (7) | Right Insula | 4.93 (28, 16, -17) |
| Cluster #12 (5) | Right Inferior Frontal Gyrus | 4.96 (25, 13, -20) |

***Partner First ∩ Stranger First***

| | | |
|---|---|---|
| Cluster #1 (2893) | Left Precentral Gyrus | 13.98 (-29, -14, 67) |
| | Left Supramarginal Gyrus | 12.48 (-59, -23, 43) |
| | Left Supramarginal Gyrus | 12.00 (-53, -32, 43) |
| | Left Superior Parietal Lobule | 11.83 (-32, -44, 61) |
| | Left Postcentral Gyrus | 11.28 (-41, -35, 70) |
| | Left Postcentral Gyrus | 10.45 (-53, -29, 58) |
| | Left Postcentral Gyrus | 10.21 (-41, -32, 49) |
| | Left Superior Parietal Lobule | 10.05 (-26, -47, 73) |
| | Left Parietal Operculum | 9.45 (-44, -5, 7) |
| | Left Supplementary Motor Area | 8.55 (-5, -5, 58) |
| Cluster #2 (803) | Right Precentral Gyrus | 6.70 (55, 7, 34) |
| | Right Inferior Frontal Gyrus | 6.47 (58, 13, 22) |
| | Right Insula | 6.24 (40, 25, 1) |
| | Right Inferior Frontal Gyrus | 6.23 (52, 16, 4) |

| | | |
|---|---|---|
| | Right Insula | 5.79 (34, 10, -11) |
| | Right Inferior Frontal Gyrus | 5.63 (34, 4, 28) |
| | Right Insula | 5.48 (40, 1, 10) |
| | Right Temporal Pole | 5.44 (31, 16, -26) |
| | Right Parahippocampal Gyrus | 5.00 (22, 1, -23) |
| Cluster #3 (411) | Right Supramarginal Gyrus | 7.00 (67, -26, 19) |
| | Right Parietal Operculum | 7.00 (55, -20, 22) |
| | Right Supramarginal Gyrus | 5.91 (37, -35, 43) |
| | Right Postcentral Gyrus | 5.90 (61, -17, 43) |
| | Right Supramarginal Gyrus | 5.73 (49, -26, 34) |
| | Right Inferior Parietal Lobule | 5.44 (37, -41, 55) |
| Cluster #4 (264) | Right Cerebellum | 9.78 (28, -56, -35) |
| | Right Cerebellum | 4.93 (10, -59, -11) |
| Cluster #5 (213) | Right Cerebellum | 9.69 (22, -56, -53) |
| | Right Cerebellum | 8.84 (16, -68, -50) |
| Cluster #6 (176) | Left Cerebellum | 6.66 (-23, -71, -53) |
| | Left Cerebellum | 6.60 (-32, -62, -56) |
| Cluster #7 (113) | Right Postcentral Gyrus | **-5.82** (28, -26, 61) |
| | Right Precentral Gyrus | **-5.55** (34, -20, 49) |
| Cluster #8 (35) | Right Caudate | 4.38 (16, 13, 10) |
| Cluster #9 (28) | Left Temporal Pole | 4.90 (-29, 7, -20) |
| Cluster #10 (14) | Right Caudate Nucleus | 4.83 (7, 1, -5) |

***Partner Second ∩ Stranger Second***

| | | |
|---|---|---|
| Cluster #1 (3121) | Left Postcentral Gyrus | 12.58 (-50, -32, 52) |
| | Left Postcentral Gyrus | 11.26 (-38, -32, 46) |
| | Left Supramarginal Gyrus | 11.04 (-56, -23, 43) |
| | Left Supramarginal Gyrus | 9.55 (-50, -35, 25) |

| | | |
|---|---|---|
| | Left Supramarginal Gyrus | 9.23 (-44, -26, 25) |
| | Left Supramarginal Gyrus | 9.17 (-56, -26, 19) |
| | Left Supramarginal Gyrus | 8.29 (-71, -29, 25) |
| | Left Postcentral Gyrus | 8.11 (-23, -44, 76) |
| | Left Precentral Gyrus | 8.11 (-32, -14, 61) |
| | Left Parietal Operculum | 7.94 (-44, -5, 10) |
| Cluster #2 (615) | Right Insula | 6.25 (43, -2, 1) |
| | Right Inferior Frontal Gyrus | 6.20 (46, 10, 19) |
| | Right Inferior Frontal Gyrus | 6.07 (55, 10, 28) |
| | Right Insula | 5.48 (25, 16, -17) |
| | Right Insula | 5.46 (40, 25, -5) |
| | Right Inferior Frontal Gyrus | 5.31 (58, 13, 7) |
| | Right Amygdala | 5.29 (31, 4, -20) |
| | Right Inferior Frontal Gyrus | 5.18 (52, 13, -2) |
| | Right Insula | 5.04 (40, 1, 16) |
| Cluster #3 (460) | Right Supramarginal Gyrus | 7.07 (58, -32, 25) |
| | Right Supramarginal Gyrus | 6.51 (58, -20, 22) |
| | Right Supramarginal Gyrus | 6.14 (52, -26, 34) |
| | Right Postcentral Gyrus | 6.05 (34, -38, 49) |
| Cluster #4 (138) | Right Cerebellum | 8.22 (25, -50, 29) |
| Cluster #5 (132) | Right Cerebellum | 8.00 (16, -65, -53) |
| Cluster #6 (86) | Right Precentral Gyrus | **-6.17** (31, -26, 61) |
| Cluster #7 (38) | Left Caudate | 5.01 (-11, 13, -2) |
| | Left Caudate | 4.68 (-20, 22, 4) |
| Cluster #8 (4) | Left Putamen | 4.57 (-17, 10, -2) |

***Partner First ∩ Stranger Second***

| | | |
|---|---|---|
| Cluster #1 (4187) | Left Precentral Gyrus | 16.16 (-29, -14, 67) |
| | Left Supramarginal Gyrus | 15.80 (-56, -26, 43) |

| | | |
|---|---|---|
| | Left Postcentral Gyrus | 14.83 (-41, -35, 67) |
| | Left Postcentral Gyrus | 14.68 (-38, -32, 46) |
| | Left Supramarginal Gyrus | 12.85 (-50, -35, 25) |
| | Left Parietal Operculum | 11.46 (-44, -5, 7) |
| | Left Superior Parietal Lobule | 11.41 (-26, -47, 73) |
| | Left Supplementary Motor Area | 9.82 (-5, -5, 58) |
| | Left Supplementary Motor Area | 9.55 (-14, 1, 67) |
| | Left Supplementary Motor Area | 9.17 (4, 7, 55) |
| Cluster #2 (2350) | Right Superior Temporal Gyrus | 8.22 (58, -35, 22) |
| | Right Parietal Operculum | 7.83 (55, -23, 22) |
| | Right Supramarginal Gyrus | 7.31 (34, -38, 46) |
| | Right Precentral Gyrus | 7.29 (52, 7, 37) |
| | Right Superior Parietal Lobule | 7.13 (34, -47, 58) |
| | Right Insula | 6.89 (34, 10, -11) |
| | Right Inferior Frontal Gyrus | 6.87 (46, 10, 19) |
| | Right Insula | 6.68 (40, 1, 16) |
| | Right Supramarginal Gyrus | 6.50 (52, -26, 34) |
| | Right Superior Frontal Gyrus | 6.49 (34, -8, 58) |
| Cluster #3 (417) | Right Cerebellum | 10.60 (25, -50, 26) |
| | Right Cerebellum | 6.25 (10, -59, -11) |
| | Right Fusiform Gyrus | 5.22 (43, -41, -17) |
| Cluster #4 (312) | Right Cerebellum | 10.62 (16, -65, -53) |
| Cluster #5 (215) | Right Middle Temporal Gyrus | 6.58 (52, -62, 1) |
| Cluster #6 (208) | Left Cerebellum | 7.81 (-23, -74, -56) |
| Cluster #7 (193) | Left Middle Temporal Gyrus | 8.66 (-56, -77, 4) |
| | Left Middle Temporal Gyrus | 7.71 (-44, -62, 7) |
| Cluster #8 (108) | Right Postcentral Gyrus | **-6.35** (28, -26, 58) |
| | Right Precentral Gyrus | **-5.36** (34, -20, 49) |
| Cluster #9 (73) | Left Cerebellum | 5.05 (-26, -65, -26) |
| Cluster #10 (47) | Left Insula | 5.26 (-32, 10, -17) |

***Stranger First ∩ Partner Second***

| | | |
|---|---|---|
| Cluster #1 (2460) | Left Postcentral Gyrus | 10.41 (-50, -32, 58) |
| | Left Rolandic Operculum | 9.55 (-47, -23, 22) |
| | Left Supramarginal Gyrus | 9.12 (-53, -32, 25) |
| | Left Supramarginal Gyrus | 8.93 (-59, -23, 43) |
| | Left Superior Parietal Lobule | 8.88 (-32, -44, 61) |
| | Left Postcentral Gyrus | 7.61 (-35, -32, 73) |
| | Left Parietal Operculum | 7.35 (-53, -2, 7) |
| | Left Postcentral Gyrus | 7.27 (-23, -44, 76) |
| | Left Supplementary Motor Area | 6.90 (-11, -5, 55) |
| Cluster #2 (387) | Right Insula | 6.22 (40, 19, -5) |
| | Right Insula | 5.68 (43, 1, -2) |
| | Right Inferior Frontal Gyrus | 5.64 (55, 16, 7) |
| | Right Inferior Frontal Gyrus | 5.29 (58, 13, 25) |
| Cluster #3 (332) | Right Supramarginal Gyrus | 6.51 (61, -29, 28) |
| | Right Supramarginal Gyrus | 5.89 (58, -17, 22) |
| | Right Supramarginal Gyrus | 5.30 (52, -26, 34) |
| | Right Inferior Parietal Lobule | 4.65 (40, -35, 46) |
| Cluster #4 (147) | Right Precentral Gyrus | **-6.08** (34, -26, 61) |
| | Right Precentral Gyrus | **-5.81** (37, -20, 52) |
| Cluster #5 (122) | Right Cerebellum | 7.07 (25, -50, 26) |
| | Right Cerebellum | 4.35 (10, -56, -14 |
| Cluster #6 (121) | Right Cerebellum | 6.77 (22, -59, -53) |
| Cluster #7 (81) | Right Supplementary Motor Area | 5.05 (7, 13, 55) |
| Cluster #8 (29) | Right Caudate | 5.73 (7, 4 -2) |
| Cluster #9 (29) | Left Caudate | 4.74 (-8, 16, -2) |
| | Left Caudate | 4.40 (-17, 22, 4) |
| Cluster #10 (23) | Right Anterior Cingulate Cortex | 4.36 (4, 19, 25) |

| | | |
|---|---|---|
| | Right Middle Cingulate Cortex | 3.96 (4, 25, 34) |
| Cluster #11 (19) | Right Insula | 4.55 (28, 22, -17) |
| Cluster #12 (14) | Right Amygdala | 4.91 (22, 1, -20) |
| | Right Amygdala | 4.42 (34, 1, -23) |
| Cluster #13 (11) | Left Middle Cingulate Cortex | 4.06 (-5, 16, 37) |
| Cluster #14 (6) | Right Caudate | 4.06 (10, 16, 1) |
| Cluster #15 (2) | Right Caudate | 5.17 (7, 1, -5) |

***Partner First ∩ Partner Second***

| | | |
|---|---|---|
| Cluster #1 (3029) | Left Precentral Gyrus | 13.90 (-29, -14, 67) |
| | Left Postcentral Gyrus | 11.66 (-32, -41, 61) |
| | Left Supramarginal Gyrus | 11.44 (-56, -26, 43) |
| | Left Postcentral Gyrus | 11.11 (-41, -35, 70) |
| | Left Parietal Operculum | 11.09 (-47, -29, 19) |
| | Left Postcentral Gyrus | 10.42 (-53, -32, 61) |
| | Left Postcentral Gyrus | 10.35 (-41, -32, 49) |
| | Left Superior Parietal Lobule | 9.93 (-23, -44, 73) |
| | Left Insula | 9.85 (-44, -2, 7) |
| | Right Supplementary Motor Area | 8.95 (4, 7, 55) |
| Cluster #2 (500) | Right Inferior Frontal Gyrus | 6.52 (58, 13, 22) |
| | Right Inferior Frontal Gyrus | 6.07 (61, 10, 10) |
| | Right Insula | 6.03 (37, 1, 16) |
| | Right Inferior Frontal Gyrus | 5.63 (40, 31, 1) |
| | Right Insula | 5.62 (43, 1, 1) |
| | Right Inferior Frontal Gyrus | 5.22 (46, 16, 1) |
| Cluster #3 (467) | Right Parietal Operculum | 7.16 (55, -20, 19) |
| | Right Supramarginal Gyrus | 5.67 (37, -35, 46) |
| Cluster #4 (136) | Right Cerebellum | 9.15 (22, -59, -50) |
| | Right Cerebellum | 8.07 (13, -68, -50) |

| | | |
|---|---|---|
| Cluster #5 (134) | Right Cerebellum | 8.58 (25, -53, -26) |
| | Right Cerebellum | 5.12 (10, -59, -11) |
| Cluster #6 (105) | Right Precentral Gyrus | **-6.20** (28, -23, 58) |
| | Right Precentral Gyrus | **-5.90** (34, -20, 49) |
| Cluster #7 (39) | Right Caudate | 4.89 (10, 10, -2) |
| | Right Caudate | 4.19 (19, 22, 4) |
| Cluster #8 (34) | Right Amygdala | 4.98 (25, 10, -17) |
| Cluster #9 (16) | Right Caudate | 5.33 (7, 4, -2) |
| Cluster #10 (2) | Right Insula | 4.22 (40, -14, 10) |

***Stranger First ∩ Stranger Second***

| | | |
|---|---|---|
| Cluster #1 (3270) | Left Postcentral Gyrus | 13.01 (-50, -32, 52) |
| | Left Supramarginal Gyrus | 11.97 (-56, -23, 43) |
| | Left Postcentral Gyrus | 11.27 (-38, -32, 46) |
| | Left Supramarginal Gyrus | 10.87 (-53, -32, 25) |
| | Left Postcentral Gyrus | 10.08 (-38, -41, 61) |
| | Left Postcentral Gyrus | 8.46 (-23, -44, 76) |
| | Left Precentral Gyrus | 8.41 (-59, 7, 31) |
| | Left Precentral Gyrus | 8.20 (-32, -14, 61) |
| | Left Parietal Operculum | 7.50 (-47, -5, 10) |
| | Left Parietal Operculum | 6.96 (-56, 4, 13) |
| Cluster #2 (952) | Right Precentral Gyrus | 7.28 (52, 7, 37) |
| | Right Precentral Gyrus | 6.87 (43, 4, 31) |
| | Right Inferior Frontal Gyrus | 6.38 (52, 19, 7) |
| | Right Amygdala/Hippocampus | 5.95 (16, -8, 14) |
| | Right Insula | 5.82 (34, 7, -14) |
| | Right Putamen | 5.61 (28, 25, 1) |
| Cluster #3 (411) | Right Supramarginal Gyrus | 6.72 (58, -32, 25) |
| | Right Supramarginal Gyrus | 6.56 (61, -17, 25) |

| | | |
|---|---|---|
| | Right Postcentral Gyrus | 6.26 (37, -38, 52) |
| | Right Postcentral Gyrus | 5.91 (58, -17, 43) |
| | Right Supramarginal Gyrus | 5.69 (52, -26, 34) |
| Cluster #4 (247) | Right Cerebellum | 9.52 (25, -50, -26) |
| | Right Cerebellum | 5.05 (7, -62, -11) |
| Cluster #5 (202) | Right Cerebellum | 8.82 (16, -65, -53) |
| Cluster #6 (135) | Left Cerebellum | 6.73 (-26, -68, -53) |
| Cluster #7 (92) | Right Postcentral Gyrus | **-5.91** (28, -26, 61) |
| Cluster #8 (5) | Left Caudate | 4.15 (-17, 19, 4) |
| Cluster #9 (3) | Left Caudate | 4.14 (-11, 16, -2) |
| Cluster #10 (3) | Left Caudate | 4.57 (-20, 22, 4) |
| Cluster #11 (2) | Left Temporal Pole | 4.10 (-47, 13, -14) |

**Table S6.** Three-dimensional multivariate modeling with factors toucher (partner, stranger) and touch location (arm, palm). All contrasts thresholded at P<0.002, cluster-size thresholded at alpha=0.05 FWE for n = 35 complete functional datasets. For each cluster under each contrast heading, size, location, maximum T, and MNI coordinates (x, y, z) are given. * = Not surviving cluster-threshold correction.

| Cluster number (size) | Peak location | T (x, y, z) |
|---|---|---|
| ***Palm > Arm*** | | |
| Cluster #1 (1793) | Left Postcentral Gyrus | 11.37 (-50, -29, 61) |
| | Left Postcentral Gyrus | 10.46 (-47, -26, 49) |
| | Left Precentral Gyrus | 9.50 (-35, -17, 64) |
| | Left Supplementary Motor Area | 9.13 (-11, -2, 52) |
| | Right Precentral Gyrus | 7.46 (37, -11, 67) |
| | Right Supplementary Motor Area | 5.71 (10, 4, 49) |
| | Left Supplementary Motor Area | 5.16 (-11, -20, 49) |
| | Left Superior Parietal Lobule | 5.05 (-26, -62, 70) |
| | Right Supplementary Motor Area | 4.90 (13, 13, 64) |
| | Left Superior Frontal Gyrus | 4.88 (-20, -2, 58) |
| Cluster #2 (503) | Right Postcentral Gyrus | 9.84 (58, -17, 52) |
| | Right Postcentral Gyrus | 9.07 (46, -23, 46) |
| | Right Postcentral Gyrus | 9.00 (49, -32, 61) |
| | Right Parietal Operculum | 4.19 (46, -20, 19) |
| | Right Supramarginal Gyrus | 3.78 (67, -20, 25) |
| Cluster #3 (454) | Right Cerebellum | 10.06 (16, -53, 23) |

| | | |
|---|---|---|
| Cluster #4 (152) | Right Cerebellum | 7.85 (16, -62, -47) |
| Cluster #5 (63) | Right Middle Frontal Gyrus | 4.66 (31, 40, 22) |
| | Right Superior Frontal Gyrus | 4.16 (22, 55, 19) |
| Cluster #6 (62) | Left Cerebellum | 4.82 (-26, -50, -32) |

***Arm > Palm***

| | | |
|---|---|---|
| Cluster #1 (128) | Left Superior Temporal Gyrus | 4.69 (-59, -50, 16) |
| | Left Superior Temporal Gyrus | 4.36 (-47, -41, 10) |
| Cluster #2 (85) | Left Postcentral Gyrus | 5.36 (-20, -38, 67) |
| | Left Precuneus | 4.94 (-17, -41, 55) |
| Cluster #3 (35) | Left Supramarginal Gyrus* | 4.63 (-44, -38, 25) |

***Stranger > Partner***

| | | |
|---|---|---|
| Cluster #1 (203) | Right Middle Frontal Gyrus | 4.84 (4, 34, 34) |
| | Left Supplementary Motor Area | 3.76 (-8, 25, 46) |
| Cluster #2 (171) | Left Inferior Frontal Gyrus | 4.76 (-47, 49, 7) |

***Interaction Site x Stroker***

| | | |
|---|---|---|
| Cluster #1 (177) | Left Postcentral Gyrus | 26.78 (-50, -35, 55) |
| | Left Precentral Gyrus | 23.22 (-44, -14, 64) |

**Table S7.** Paired T-tests for partner vs stranger during each of 2 functional runs (first, second), modeled with linear mixed effects and weighted by individual mean cortisol levels as covariate. All contrasts thresholded at P<0.002, cluster-size thresholded at alpha=0.05 FWE for n = 35 complete functional datasets. For each cluster under each contrast heading, size, location, maximum T, and MNI coordinates (x, y, z) are given.

| Cluster number (size) | Peak location | T (x, y, z) |
|---|---|---|
| ***Partner First > Stranger Second*** | | |
| Cluster #1 (495) | Left Anterior Cingulate Cortex | 7.94 (-8, 25, 16) |
| | Left Anterior Cingulate Cortex | 7.14 (-2, 34, 16) |
| | Left Anterior Cingulate Cortex | 7.08 (-11, 46, 10) |
| | Right Superior Medial Gyrus | 5.95 (7, 58, 10) |
| | Right Superior Medial Gyrus | 5.49 (-8, 64, 13) |
| | Left Mid Orbitofrontal Gyrus | 5.05 (-8, 43, -5) |
| | Right Mid Orbitofrontal Gyrus | 4.70 (4, 67, -8) |
| | Left Mid Orbitofrontal Gyrus | 4.43 (-8, 52, -11) |
| Cluster #2 (180) | Right Calcarine Gyrus | 6.11 (25, -74, 7) |
| | Right Lingual Gyrus | 5.54 (7, -71, -2) |
| | Right Calcarine Gyrus | 5.54 (10, -74, 10) |
| | Right Calcarine Gyrus | 5.42 (-11, -65, 4) |
| | Left Calcarine Gyrus | 4.97 (-2, -80, 10) |
| Cluster #3 (81) | Left Temporal Pole | 6.32 (-59, 7, -5) |
| | Left Paretal Operculum | 6.32 (-59, -2, 13) |
